# Alternative splicing regulates the physiological adaptation of the mouse hind limb postural and phasic muscles to microgravity

**DOI:** 10.1101/2021.05.25.445491

**Authors:** Mason Henrich, Pin Ha, John S. Adams, Chia Soo, Kang Ting, Louis Stodieck, Rene Chun

## Abstract

Muscle atrophy and fiber type alterations are well-characterized physiological adaptations to microgravity with both understood to be primarily regulated by differential gene expression (DGE). While microgravity-induced DGE has been extensively investigated, adaptations to microgravity due to alternative splicing (AS) have not been studied in a mammalian model. We sought to comprehensively elucidate the transcriptomic underpinnings of microgravity-induced muscle phenotypes in mice by evaluating both DGE and changes in AS due to extended spaceflight. Tissue sections and total RNA were isolated from the gastrocnemius and quadriceps, postural and phasic muscles of the hind limb, respectively, of 32-week-old female BALB/c mice exposed to microgravity or ground control conditions for nine weeks. Immunohistochemistry disclosed muscle type-specific physiological adaptations to microgravity that included i) a pronounced reduction in muscle fiber cross-sectional area in both muscles and ii) a prominent slow-to-fast fiber type transition in the gastrocnemius. RNA sequencing revealed that DGE and AS varied across postural and phasic muscle types with preferential employment of DGE in the gastrocnemius and AS in the quadriceps. Gene ontology analysis indicated that DGE and AS regulate distinct molecular processes. Various non-differentially expressed transcripts encoding musculoskeletal proteins (*Tnnt3, Tnnt1, Neb, Ryr1,* and *Ttn*) and muscle-specific RNA binding splicing regulators (*Mbnl1 and Rbfox1*) were found to have significant changes in AS that altered critical functional domains of their protein products. In striking contrast, microgravity-induced differentially expressed genes were associated with lipid metabolism and mitochondrial function. Our work serves as the first comprehensive investigation of coordinate changes in DGE and AS in large limb muscles across spaceflight. We propose that substantial remodeling of pre-mRNA by AS is a major component of transcriptomic adaptation of skeletal muscle to microgravity. The alternatively spliced genes identified here could be targeted by small molecule splicing regulator therapies to address microgravity-induced changes in muscle during spaceflight.

## Introduction

Prolonged disuse of skeletal muscle in microgravity environments, often referred to as mechanical unloading, precipitates acute muscle atrophy characterized by a loss of muscle mass and strength (Adams et al., 1985). Similar microgravity-associated muscle phenotypes have been identified in the limb muscles of mice (Sandonà et al., 2012), rats (Martin et al., 1985), and monkeys (Shenkman et al., 1994) exposed to microgravity as well as sedentary human populations such as the elderly (Suetta et al., 2012) and handicapped (Giangregorio et al., 2006). While countermeasures, chief among them being regular exercise, are beneficial (Gao et al., 2018), these interventions fail to completely prevent microgravity-induced atrophy (Buckley, 2006). Ultimately, extended spaceflight results in a 4.6 to 8.8% reduction in muscle size and a reciprocal 5.9 to 8.8% increase in fat content (Burkhart et al., 2019).

While aggressive muscle conditioning and rehabilitation on Earth eventually restore muscle mass and strength, this process can take anywhere from a few months to four years to occur (Burkhart et al., 2019), with differences in rehabilitation time attributed to functional distinctions between postural and phasic muscle types. Postural muscles are composed primarily of slow twitch muscle fibers and phasic muscles are composed primarily of fast twitch muscle fibers, a distinction that underlines their respective functions (Burkholder et al., 1994). Although both are required for locomotion, postural muscles (i.e gastrocnemius) act primarily to support the body’s posture in Earth’s gravitational field, and phasic muscles (i.e. quadriceps) act primarily to support explosive movement (Charles et al., 2016). Attributable to their contrasting roles in motor function, postural and phasic muscles respond differently to extended spaceflight. Most importantly, exposure to microgravity significantly reduces the requirement for the postural component of motor function, a perturbation that precipitates an adaptive migration from a primarily slow twitch to a primarily fast twitch fiber type composition (Shenkman, 2016). The prominent slow-to-fast fiber type transition in postural muscles is speculated to account for their resistance to microgravity-induced atrophy (Fitts et al., 2001), as while phasic muscles undergo “typical” atrophic changes to microgravity, including loss of muscle mass and decreased strength of contraction (Roy et al., 1996; Recktenwald et al., 1999), postural muscles are not as affected by microgravity, displaying minimal loss of muscle mass and, in some cases, increased amplitude of muscle contraction (Roy et al., 1996; Recktenwald et al., 1999).

The development of high-throughput sequencing technologies has spurred extensive interest in elucidating the transcriptomic underpinnings of muscle type-specific microgravity-induced phenotypes. Most interest has been paid to the differential gene expression (DGE) of myosin heavy chain genes, long characterized as the canonical mechanism of fiber type transitions. There is evidence in humans (Trappe et al., 2009) as well as mice (Sandonà et al., 2012), rats (Martin et al., 1985), and monkeys (Shenkman et al., 1994) to support microgravity-induced downregulation of the myosin heavy chain gene *MYHC7*, which encodes the slow twitch muscle fiber protein MYHC I and upregulation of the myosin heavy chain genes *MYH2, MYH1,* and *MYH4* which encode fast twitch muscle fiber proteins MYHC IIA, MYHC IID/X, and MYHC IIB, respectively. In addition, microgravity-induced DGE has been shown to regulate gene networks related to protein, lipid, and carbohydrate metabolism as well as mitochondrial dysfunction in skeletal muscle (Chakraborty et al., 2020). However, a focus restricted to the level of gene expression fails to fully characterize the amplifying actions of alternative splicing (AS) on the host’s transcriptome; for example, AS can affect a multi-fold increase in the diversity translatable mRNA isoforms over what can be accounted for by the 18,000 genes in the human genome. This additional complexity is attributable to the process of RNA binding protein (RBP)-mediated AS (Nilsen and Graveley, 2010). Approximately 90% of human genes have been found to exhibit alternatively spliced isoforms (Wang et al., 2008). While there are multiple types of AS events, the most prevalent and best characterized AS types are skipped exon (SE) and mutually exclusive exon (MXE) events (Park et al., 2018). Considering the well-annotated role of AS in musculoskeletal development (Nikanova and Kao, 2020; Nakka et al., 2018; Brinegar et al., 2017), muscle fiber type composition (Briggs Schachat, 1996; Tang et al., 2015; Lam et al., 2018), and atrophic musculoskeletal diseases (Kimura et al., 2005; Tang et al., 2015; Lam et al., 2018), there has been only a single examination of microgravity-induced AS conducted, and that study focused on *Arabidopsis thaliana*, a species of flowering plant (Beisel et al., 2019).

Here we set out to characterize, for the first time in an animal model, the regulatory role of AS in the physiological adaptation to extended spaceflight. Tissue sections and total RNA were isolated from the gastrocnemius and quadriceps muscles of 32-week-old female BALB/c mice exposed to either microgravity or ground control conditions for a total of nine weeks. Using immunohistochemistry and RNA sequencing (RNA-seq), we aimed to i) annotate muscle-type-specific patterns of muscle atrophy and fiber type changes as well as ii) characterize functionally significant AS changes in non-differentially expressed striated muscle genes that describe a previously uninvestigated means of modifying the transcriptomic response to microgravity.

## Results

### Fiber type composition and size of postural and phasic muscles are differentially perturbed by spaceflight

Compared to ground controls, the overall abundance of slow-twitch fibers in the gastrocnemius decreased from 18% to 5%, with a reciprocal increase in the abundance of fast twitch fibers from 80% to 96% following nine weeks of microgravity (Figure 1A, 1C). In addition, slow and fast twitch myosin heavy chains were co-expressed at the level of individual muscle fibers (Figure 1B). By contrast, there was no evidence of a slow-to-fast fiber type transition in the quadriceps as a consequence of a 100% fast twitch fiber type composition (Figure 1A).

**Figure 1.**
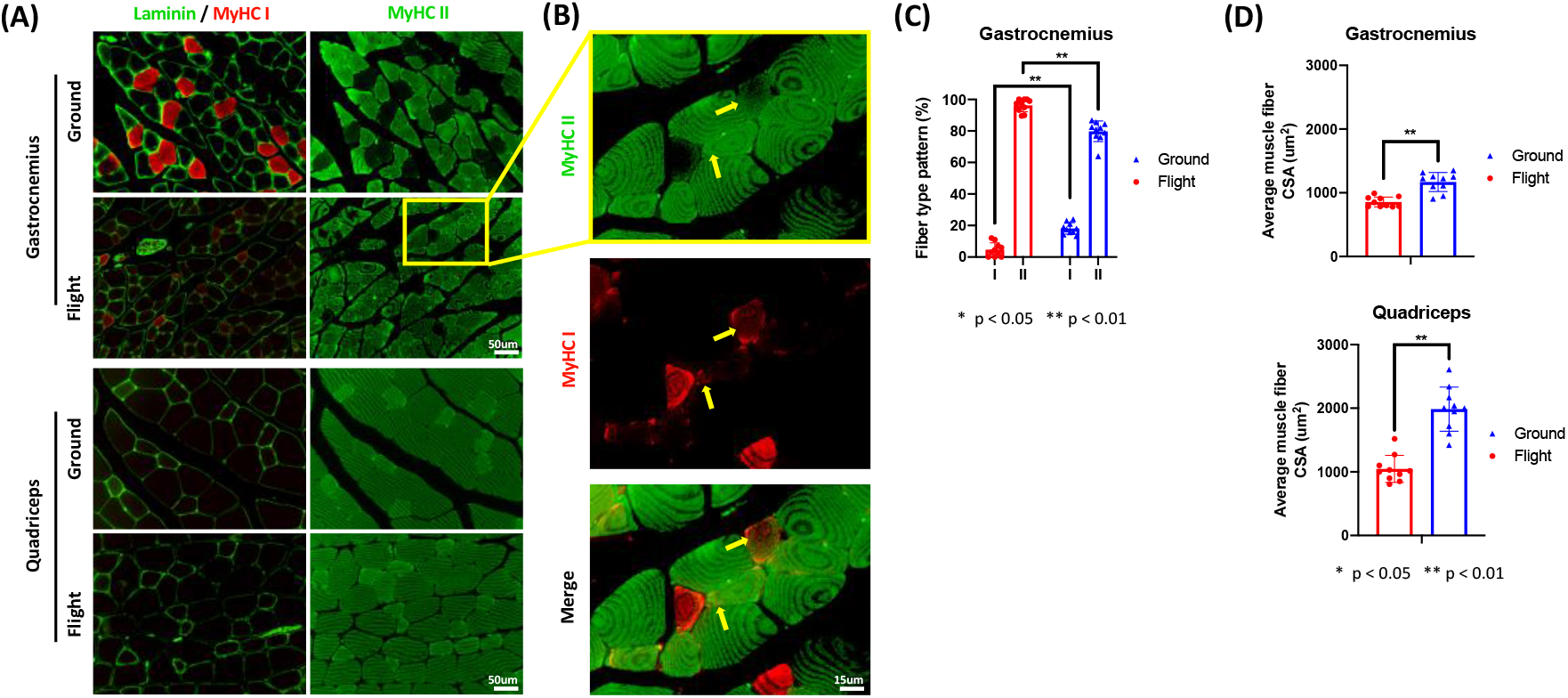
**(A)** Representative immunofluorescence images of gastrocnemius and quadriceps stained for laminin and MyHC isoforms (MyHC I: red, II: green) in flight and ground groups. **(B)** Some myofibers (yellow arrows) were co-stained by both MyHC isoforms (MyHC I: red, II: green). **(C)** Quantification of the muscle fiber type distribution of gastrocnemius (n = 10). **(D)** Quantification of the myofiber cross sectional area (CSA) in both gastrocnemius and quadriceps (n = 10 per group).

On average, the cross-sectional area of muscle fibers composing the gastrocnemius decreased from 1170 ± 141 um^2^ to 856 ± 74 um^2^, representing a 27% reduction in muscle fiber size as a result of extended exposure to microgravity. The average cross-sectional area of muscle fibers composing the quadriceps decreased from 1986 ± 330 um^2^ to 1046 ± 201 um^2^, representing a 47% reduction in muscle fiber size (Figure 1D). Therefore, the quadriceps underwent more significant microgravity-induced atrophy (approximately 1.8x more based on fiber area) than did the gastrocnemius.

### DGE and AS are muscle-type-specific and functionally distinct mechanisms of microgravity-induced transcriptome regulation

In response to extended spaceflight, the gastrocnemius and quadriceps of adult mice displayed evidence of change in both DGE and AS. However, the utilization of these two mechanisms of transcriptome regulation varied significantly between each of these muscle types. First, there was minimal overlap between microgravity-induced change in the transcriptomes of the gastrocnemius and quadriceps; only 9% of all differentially expressed genes and 6% of all alternatively spliced genes were held in common between these two muscles (Figure 2A). In addition, there was evidence of an inverse relationship between DGE and AS in the gastrocnemius and quadriceps; the ratio of differentially expressed genes to alternatively spliced genes approximated to 5:2 in the gastrocnemius and 2:5 in the quadriceps (Figure 2A).

**Figure 2.**
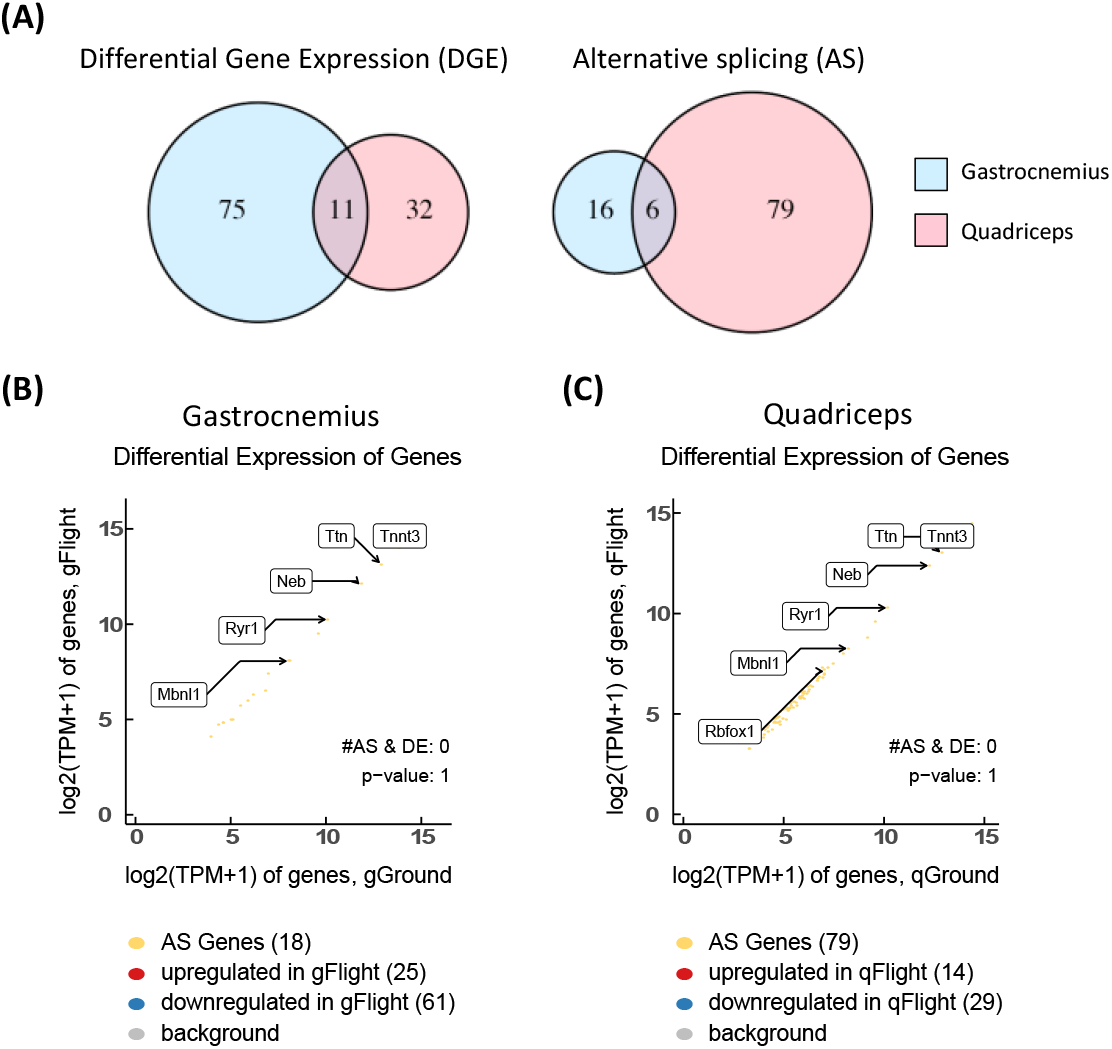
**(A)** Venn diagrams comparing differentially expressed (left) and alternatively spliced (right) genes in the gastrocnemius and quadriceps. The size of each circle is representative of the number of genes that were differentially expressed or alternatively spliced in each muscle, with blue and red circles representing such genes from the gastrocnemius and quadriceps, respectively. Overlapping regions depict genes that were held in common between the two muscles. Scatter plots comparing gene expression in ground vs. flight in the gastrocnemius **(B)** and quadriceps **(C)**. Red and blue dots represent up-regulated and down-regulated genes, respectively. Genes with significant AS changes (denoted as AS genes) are depicted in yellow. Background genes, e.g. those with no differential gene expression and no exon skipping changes, are shown in grey.

Not only were DGE and AS differentially employed in each muscle investigated, these two mechanisms of transcriptome regulation were also enriched in functionally distinct gene networks. All alternatively spliced genes were non-differentially expressed (Figure 2C), indicating that AS and DGE are differentially deployed to affect different transcripts. In addition, gene ontology analyses (Supplemental Tables 1, 2, and 3) of all differentially expressed (Supplemental Table 4) and alternatively spliced (Supplemental Tables 5 and 6) genes suggest that AS and DGE regulate unique molecular processes. Upregulated genes in the gastrocnemius quadriceps displayed insignificant and nonspecific gene network enrichment, respectively. However, downregulation of gene networks involved in mitochondrial function and lipid metabolism were found in the gastrocnemius and quadriceps, respectively. While there was some evidence to support downregulation of musculoskeletal genes in the gastrocnemius, alternatively spliced genes in both the gastrocnemius and quadriceps were overwhelmingly enriched for processes relating to the structure and function of the muscle sarcomere.

### *Troponin T3* (*Tnnt3*) and *Troponin T1* (*Tnnt1*) are alternatively spliced in favor of isoforms that enhance fast twitch muscle fiber function

Despite evidence at the level of immunostaining of a slow-to-fast fiber type transition in the gastrocnemius of adult mice exposed to spaceflight, there was no evidence at the level of RNA sequencing of differential expression of myosin heavy chain genes. In fact, the only differentially expressed genes with annotated roles in muscle fiber composition were those encoding small, regulatory proteins in the muscle sarcomere (*Tnni1, Tnnc1,* and *Myl*, Supplemental Table 4). Interestingly, *Troponin T*, which encodes a subunit of the troponin complex that facilitates muscle contraction via functional interaction with two of these differentially expressed genes (*Tnni1* and *Tnnc1*; Farah and Reinach, 1995), was not differentially expressed in either the gastrocnemius or the quadriceps (Supplemental Table 4). Instead, *Tnnt3* and *Tnnt1*, the fast- and slow-twitch isoforms of *Troponin T*, respectively, were alternatively spliced (Figure 3, 4) in favor of transcripts encoding protein isoforms that functionally support fast-twitch fiber environments.

**Figure 3.**
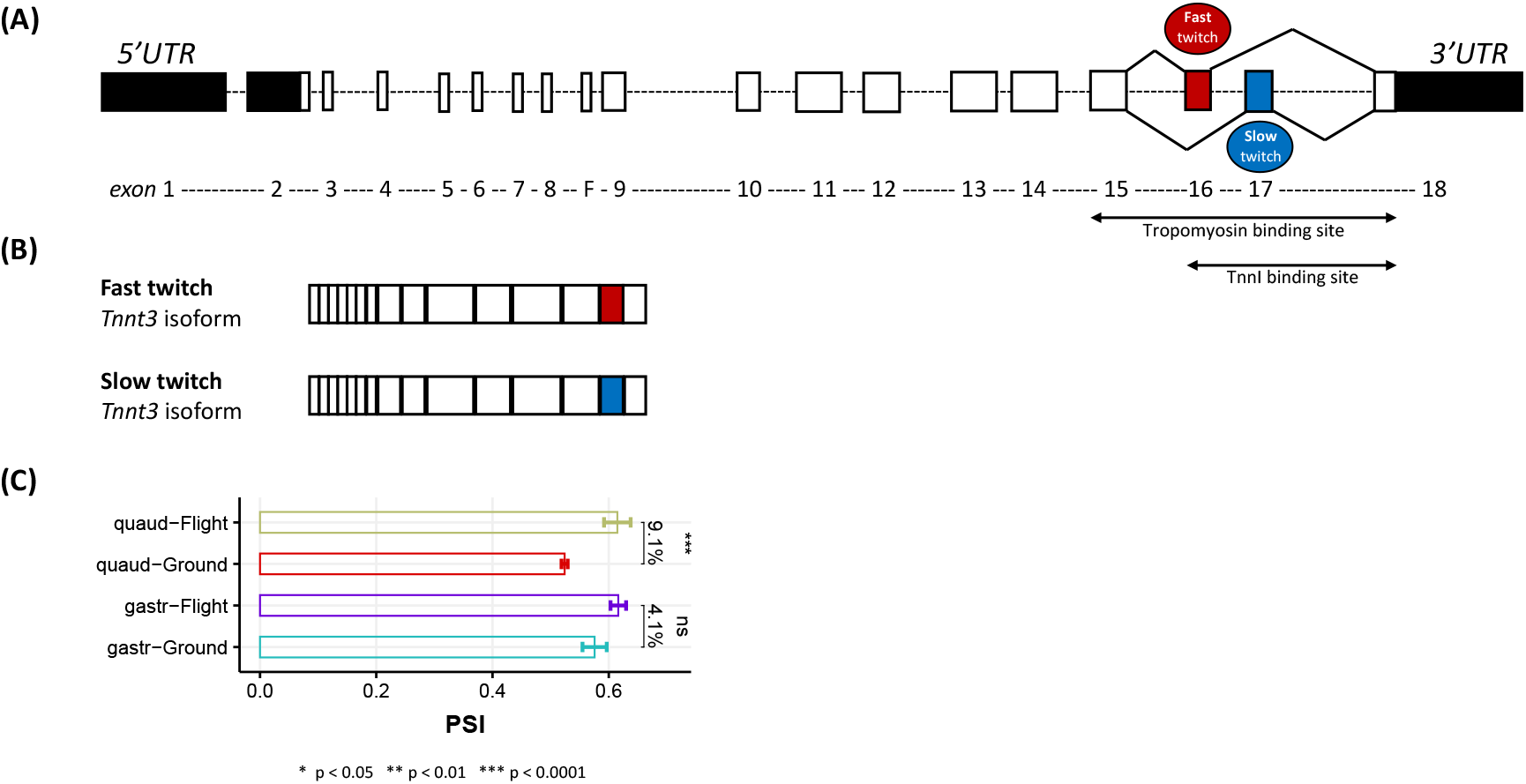
**(A)** Gene structure representation of the *Tnnt3* gene. Bars and dashed lines represent exons and introns, respectively, with exon numbering below. Untranslated regions (UTR) are denoted in black. Solid lines connecting exons 16 (depicted in red and labeled the fast twitch exon) and exon 17 (depicted in blue and labeled the slow twitch exon) to nearby splice junctions represent a mutually exclusive splicing event in a the 3’ region of the transcript that encodes for the tropomyosin and troponin I binding sites of the resulting troponin T protein. **(B)** Protein structure representation of the two *Tnnt3* isoforms. The fast twitch *Tnnt3* isoform has exon 16 included and exon 17 excluded while the slow twitch *Tnnt3* isoform has exon 16 excluded and exon 17 included. **(C)** Bar plot depicting the PSI (percent spliced in) value of the exon 16/17 mutually exclusive exon (MXE) event in ground and flight for quadriceps (top) and gastrocnemius (bottom). The delta PSI value indicates the difference in percent inclusion of exon 16/17 between flight and ground. Asterisks represent significance by two-tailed t-test.

**Figure 4.**
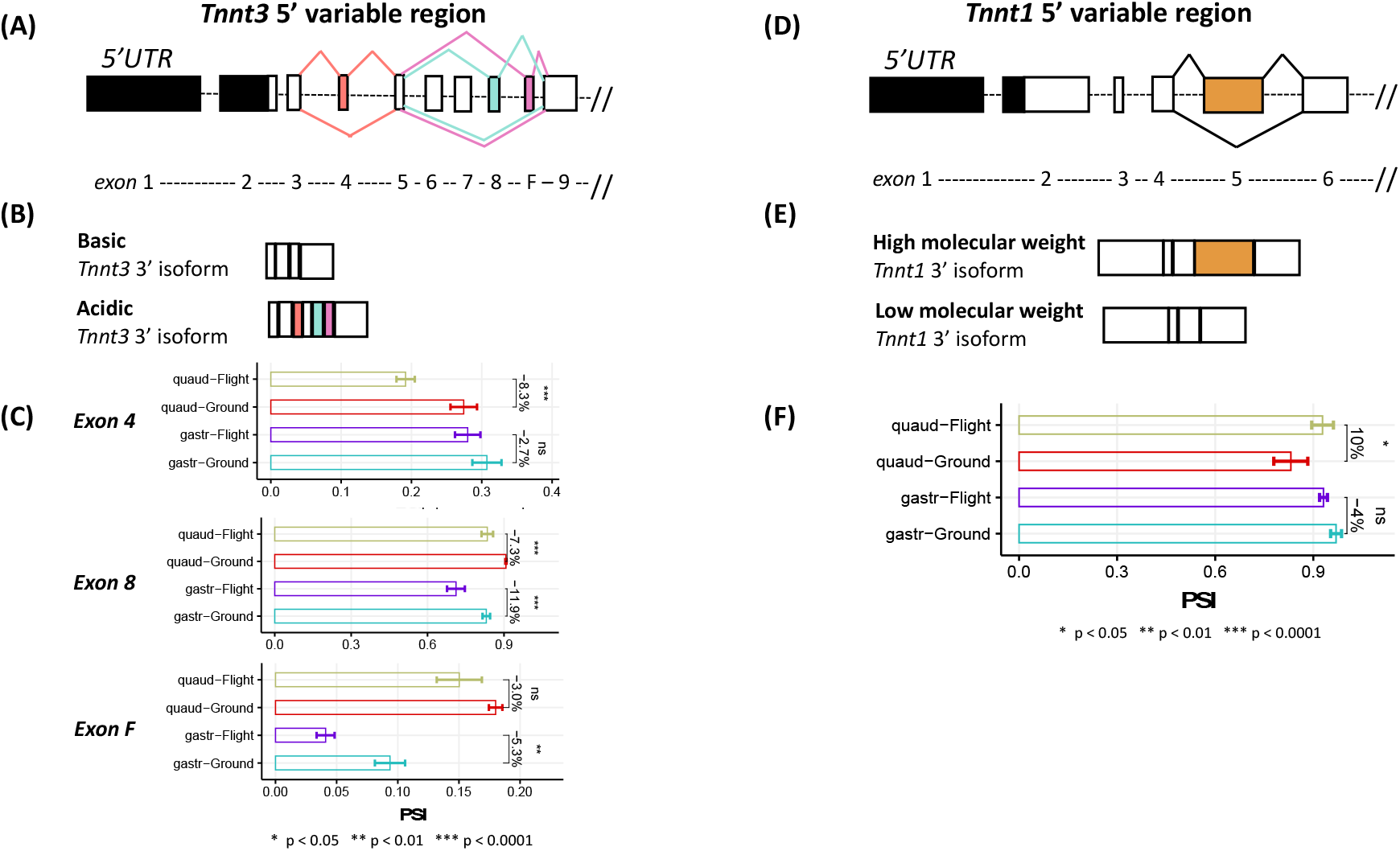
**(A)** Gene structure representation of the 5’ variable region (exons 1-9) of *Tnnt3* gene. Bars and dashed lines represent exons and introns, respectively, with exon numbering below. Untranslated regions (UTR) are denoted in black. Exon 4 (orange-red), 8 (mint), and F (pink-purple) are all alternatively spliced in an exon skipping manner with solid lines of corresponding color connecting these exons to their respective splice junctions. **(B)** Protein structure representation of two 5’ *Tnnt3* isoforms. The basic *Tnnt3* isoform has exon 4, 8, and F all excluded while the acid *Tnnt3* isoform has exon 4, 8, and F all included. **(C)** Bar plot depicting the PSI (percent spliced in) value for each exon (4, 8, and F) in ground and flight for quadriceps (top) and gastrocnemius (bottom). The delta PSI value indicates the difference in percent inclusion of each exon between flight and ground. Asterisks represent significance by two-tailed t-test. **(D)** Gene structure representation of the 5’ variable region (exons 1-6) of *Tnnt1* gene. Bars and dashed lines represent exons and introns, respectively, with exon numbering below. Untranslated regions (UTR) are denoted in black. Exon 5 (orange) is alternatively spliced in an exon skipping manner with solid lines connecting this exon to its respective splice junctions. **(E)** Protein structure representation of two 5’ *Tnnt1* isoforms. The high molecular weight *Tnnt1* isoform has exon 5 included while the low molecular weight *Tnnt1* isoform has exon 5 excluded. (**F)** Bar plot depicting the PSI (percent spliced in) value for exon 5 in ground and flight for quadriceps (top) and gastrocnemius (bottom). The delta PSI value indicates the difference in percent inclusion of exon 5 between flight and ground. Asterisks represent significance by two-tailed t-test.

The 3’ region of *Tnnt3* encodes the C-terminal binding domains for troponin I and tropomyosin (Stefancsik et al., 2003); therefore, the generation of two distinct *Tnnt3* isoforms via the mutual exclusion of two 3’ exons (Figure 3A, 3B) influences the affinity of troponin T3 for its binding partners. Previous research suggests that exon 16 and exon 17 of *Tnnt3* vary in their sequence similarity to the functionally equivalent exon in *Tnnt1*. Specifically, exon 17 shows a much higher degree of similarity (61%) than does exon 16 (32%). As a consequence the binding affinity of the exon 17-included troponin T3 isoform is higher for the slow-twitch isoforms of troponin I and tropomyosin and, reciprocally, the binding affinity of the exon 16-included Troponin T3 isoform is higher for the fast-twitch isoforms of troponin I and tropomyosin (Wang and Jin, 1997). In the quadriceps of mice exposed to spaceflight, the exon 16-included *Tnnt3* isoform was 9.1% more abundant, while the exon 17-included *Tnnt3* isoform was reciprocally 9.1% less abundant (Figure 3C). In the gastrocnemius of mice exposed to spaceflight, exon 16-included *Tnnt3* isoform was 4.1% more abundant while the exon 17-included *Tnnt3* isoform was reciprocally 4.1% less abundant, just failing to meet statistical significance (Figure 3C). Together, these results support an exon 17-to-16 transition in the quadriceps during extended spaceflight.

While the N-terminal variable region of troponin T3 has no known binding partners in the thin filament regulatory system, alternative splicing within the 5’ variable region of *Tnnt3* (Figure 4A) generates various isoforms that fall into either acidic residue-enriched or basic residue-enriched isoform classes (Figure 4B). Changes in the charge of the 5’ variable region alters the three-dimensional structure of the resulting protein (Biesiadecki et al., 2007; Wei and Jin, 2011) resulting in changes in calcium (Ca^2+^) sensitivity of the troponin complex. Specifically, basic isoforms tend to have a greater Ca^2+^ sensitivity to contraction that contributes to their preferential utilization in fast-twitch skeletal muscle fibers (Briggs et al., 1996; Ogut et al., 1999). In comparison to acidic isoforms, basic isoforms preferentially exclude exons 4, 8, and F (Figure 4B) (Wang and Jin, 1997). In the quadriceps there was significant exclusion of exons 4 (−8.3%) and 8 (−7.3%). While exon F was included 3.0% less in spaceflight conditions, this decrease was insufficient to meet statistical significance. Likewise, in the gastrocnemius there was significant exclusion of two of the exons, 8 (−11.9%) and F (−5.3%), while exon 4 was included 2.7% less in spaceflight conditions, just failing to meet statistical significance (Figure 4C). Together, these results provide evidence of an acidic-to-basic transition of the 5’ variable region of *Tnnt3* in both the quadriceps and gastrocnemius during extended spaceflight.

Similar to *Tnnt3*, *Tnnt1* undergoes AS within its 5’ variable region (Figure 4D). While the resulting isoforms do not differ in their acid or basic nature, the functional consequences of AS are comparable. The exclusion or inclusion of *Tnnt1* exon 5 creates lower and higher molecular weight protein isoforms, respectively (Figure 4E). Changes in the molecular weight of the 5’ variable region alters the three-dimensional structure of the resulting protein (Wei and Jin, 2011), similarly influencing the calcium (Ca^2+^) sensitivity of the troponin complex. Specifically, the higher molecular weight isoform (exon 5-included), tends to have a greater sensitivity to Ca^2+^, a change that contributes to its preferential utilization in fast-twitch skeletal muscle fibers (Briggs et al., 1996; Ogut et al., 1999). In the quadriceps of mice exposed to spaceflight, exon 5 was significantly more abundant (10%). Whereas in the gastrocnemius of mice exposed to spaceflight, exon 5 was 4% less abundant, just failing to reach significance (Figure 4F). Therefore, our results support a lower-to-higher molecular weight *Tnnt1* isoform transition in the quadriceps during extended spaceflight.

### The transition between distinct, developmentally immature and mature isoforms of the *nebulin* (*Neb*) and *ryanodine receptor 1* (*Ryr1*) genes during spaceflight

There is extensive evidence to suggest that developmental transitions in muscle phenotype are under the control of AS. At a transcriptomic level, the employment of AS fluctuates across development, with a surge in AS events immediately after birth and a resurgence of AS in adulthood (Brinegar et al., 2017). At an individual transcript level, certain exons are included to a greater degree in younger individuals while others to a greater degree in older individuals. This is the case of *Neb* and *Ryr1* (Donner et al., 2006; Tang et al., 2015). The *Neb* gene encodes nebulin, an actin-binding protein that is localized to the thin filament of the sarcomeres in skeletal muscle. Nebulin is a very large protein (600–900 kDa) that binds as many as 200 actin monomers during muscle contraction (Chu et al., 2016). *Neb* possesses two mutually exclusive exons (127 and 128), with the exon 127-included isoform being more abundant in developmentally immature mice and the exon 128-included isoform being more abundant in developmentally mature mice (Figure 5A) (Donner et al., 2006). Appropriately, these two isoforms are labeled as the fetal and adult *Neb* isoforms, respectively (Figure 5B). The fetal isoform is significantly more abundant in the quadriceps (38 %). Whereas in the gastrocnemius of mice exposed to spaceflight, the fetal isoform was 4% less abundant, just failing to reach significance (Figure 5C).Therefore, spaceflight-induced AS seems to induce a reversion towards a more developmentally primitive isoform of the mRNA and encoded protein in the quadriceps.

**Figure 5.**
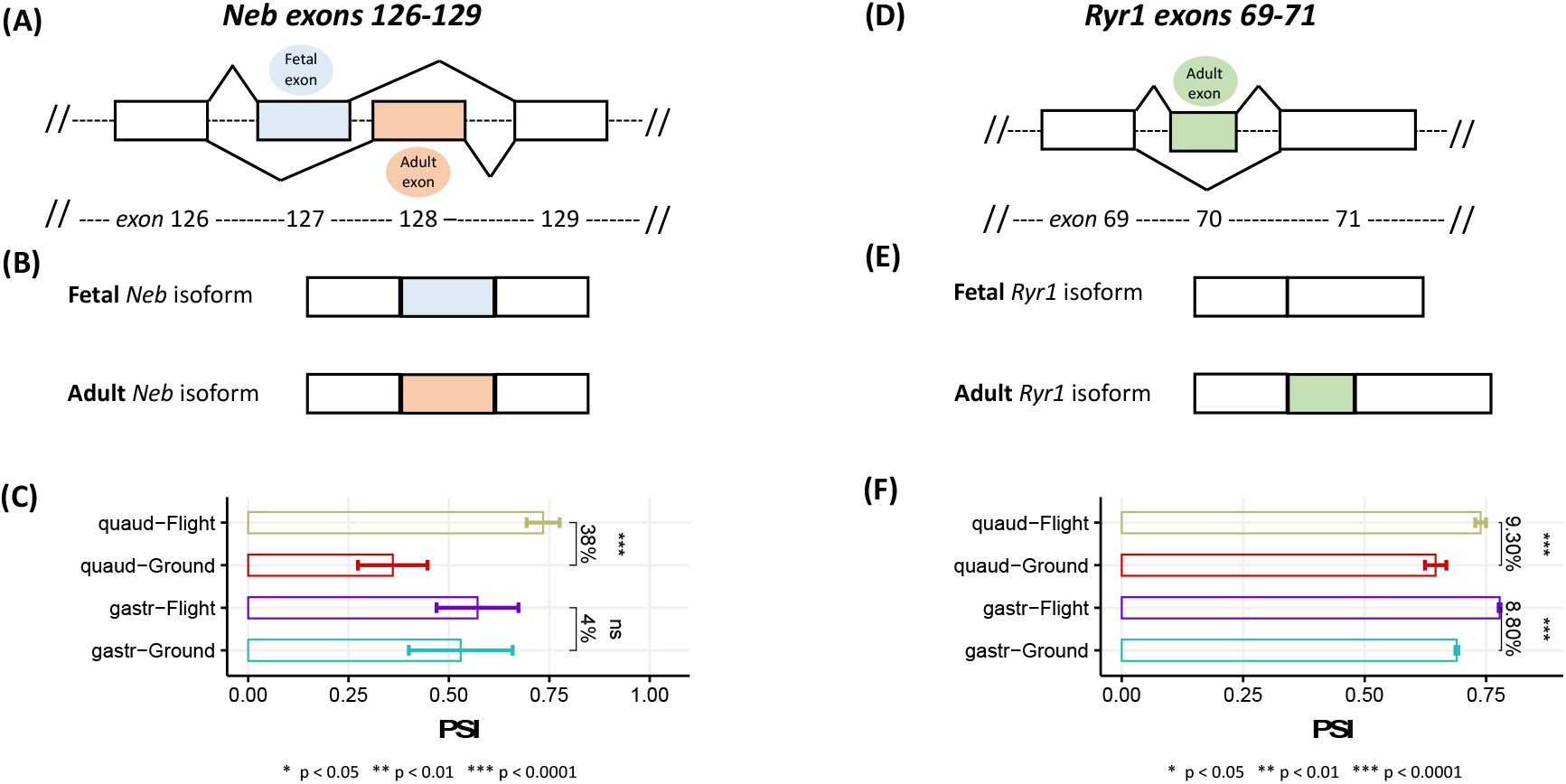
**(A)** Gene structure representation of exons 126-129 of *Neb* gene. Bars and dashed lines represent exons and introns, respectively, with exon numbering below. Solid lines connecting exons 127 (depicted in light blue and labeled the fetal exon) and exon 128 (depicted in peach and labeled the adult exon) to nearby splice junctions represent a mutually exclusive splicing. **(B)** Protein structure representation of the two *Neb* isoforms. The fetal *Neb* isoform has exon 127 included and exon 128 excluded while the adult *Neb* isoform has exon 127 excluded and exon 128 included. **(C)** Bar plot depicting the PSI (percent spliced in) value of the exon 127/128 mutually exclusive exon (MXE) event in ground and flight for quadriceps (top) and gastrocnemius (bottom). The delta PSI value indicates the difference in percent inclusion of exon 127/128 between flight and ground. Asterisks represent significance by two-tailed t-test. **(D)** Gene structure representation of exons 69-71 of Ryr 1 gene. Bars and dashed lines represent exons and introns, respectively, with exon numbering below. Exon 70 (green) is alternatively spliced in an exon skipping manner with solid lines connecting this exons to its respective splice junctions. **(E)** Protein structure representation of two *Ryr1* isoforms. The fetal *Ryr1* isoform has exon 70 excluded while the adult *Ryr1* isoform has exon 70 included. **(F)** Bar plot depicting the PSI (percent spliced in) value for exon 70 in ground and flight for quadriceps (top) and gastrocnemius (bottom). The delta PSI value indicates the difference in percent inclusion of exon 70 between flight and ground. Asterisks represent significance by two-tailed t-test.

A similar phenomenon is seen in the context of *Ryr1*, a gene that encodes a calcium release channel in skeletal muscle (Hernandez-Ochoa et al., 2016). *Ryr1* is alternatively spliced in an exon skipping manner to generate two isoforms, with the exon 70-excluded isoform being more abundant in developmentally immature mice and the exon 70-included isoform being more abundant in developmentally mature mice (Figure 5D) (Tang et al., 2015). Appropriately, these two isoforms are labeled as the fetal and adult *Ryr1* isoforms, respectively (Figure 5E). The adult isoform is significantly more abundant in both the quadriceps (9.3%) and gastrocnemius (8.8%) of adult mice exposed to spaceflight (Figure 5F). Therefore, in contrast to *Neb*, spaceflight-induced AS of *Ryr1* seems to induce a progression towards a more developmentally mature isoform of the mRNA and encoded protein in both muscles.

### Microgravity-induced AS of large musculoskeletal genes, *titin (Ttn)* and *nebulin (Neb)*, precipitates atrophic phenotypes

The splicing changes discussed so far have been distinct AS events involving exons with likely functional consequence in the encoded protein. However, numerous AS events may also act concomitantly to impact the structure and function of large musculoskeletal with significant alterations to the size of these proteins possibly precipitating the development of microgravity-induced atrophic muscle phenotypes. This appears to be the case with *Ttn* and *Neb*.

Titin, which is encoded by the gene *Ttn*, regulates the elasticity and contractile strength of the sarcomere via the length of its PEVK “spring” domain; the longer the length of the PEVK domain, the weaker the contractile strength of the sarcomere (Ottenheijm er al., 2009). In the quadriceps of mice exposed to microgravity, there was evidence of extensive AS within the PEVK domain (Figure 6A). The average ΔPSI value across all significant AS events was 8.10% (Figure 6B). When taking into account the size of these exons in relation to the domain as a whole, AS may account for up to a 2.0% increase in the size of the PEVK domain in the quadriceps. Considering that a longer PEVK domain correlates with weaker contractile strength, spaceflight-induced extension of the PEVK domain might weaken the overall contractile strength of this muscle. In the gastrocnemius of mice exposed to microgravity, there was a relatively equal distribution of positive and negative ΔPSI events among the 10 significant AS events within the PEVK domain. As such, the average ΔPSI value across all AS events was −1.97% (Figure 6C). Therefore, a lack of spaceflight-induced alterations to the PEVK domain might maintain the overall contractile strength of this muscle.

**Figure 6.**
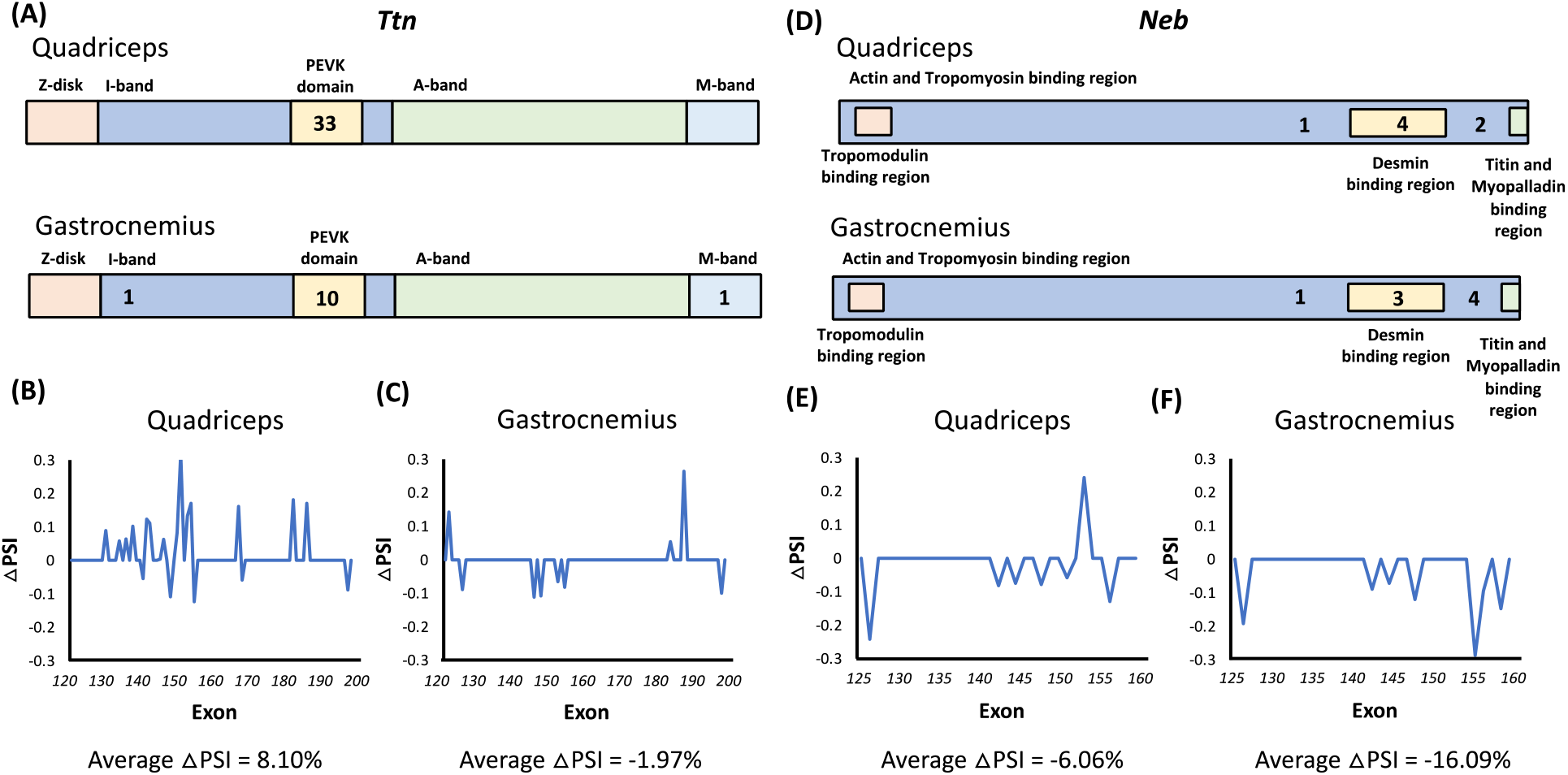
**(A)** Protein structure representation of *Ttn* gene. Colored regions represent protein domains with pink, dark blue, yellow, green, and light blue regions corresponding to Z-disk, I-band, PEVK, A-band, and M-and regions of Ttn respectively. Solid numbers represent the number of significant AS events within the corresponding region of interest in the quadricep (top) and gastrocnemius (bottom). Line graphs displaying delta PSI values for exons within the *Ttn* PEVK domain (exons 122-202) that underwent significant AS between ground and flight grounds in **(B)** quadriceps and **(C)** gastrocnemius. Positive delta PSI value events represent exons that were included more in the flight group while negative delta PSI value events represent exons that were included less in the flight group. Non-alternatively spliced and non-significantly alternatively spliced exons were provided a delta PSI value of 0. **(D)** Protein structure representation of *Neb* gene. Colored regions represent protein domains with blue, pink, yellow, and green regions corresponding to the actin and tropomyosin binding region (which spans the entire protein length), tropomodulin binding region, desmin binding region, and titin and myopalladin binding regions, respectively, in the quadricep (top) and gastrocnemius (bottom). Line graphs displaying delta PSI values for exons within the variable region of *Neb* (exons 127-159) that underwent significant AS between ground and flight grounds in **(E)** quadriceps and **(F)** gastrocnemius. Positive delta PSI value events represent exons that were included more in the flight group while negative delta PSI value events represent exons that were included less in the flight group. Non-alternatively spliced and non-significantly alternatively spliced exons were provided a delta PSI value of 0.

Nebulin, which is encoded by the gene *Neb,* regulates thin filament length. In contrast to titin, a shorter length correlates with a weaker contractile strength of the sarcomere (Bang et al., 2006). In both the quadriceps and gastrocnemius of mice exposed to microgravity, there was extensive AS within the 3’ region of the *Neb* transcript, with involved exons encoding the actin, tropomyosin, or desmin binding regions of nebulin (Figure 6D). The average ΔPSI value across all AS events was −6.06% and −16.09% in the quadriceps and gastrocnemius, respectively (Figure 6E, 6F). When taking into account the size of these exons in relation to the size of the 3’ variable region, AS may account for approximately 1.0% and 3.5% decreases in the overall size of nebulin in the quadriceps and the gastrocnemius, respectively. Considering that a shorter length correlates with weaker contractile strength, spaceflight-induced shortening of nebulin might weaken the overall contractile strength of these two muscles.

### Musculoskeletal splicing regulators,*muscleblind-like splicing regulator 1* (*Mbnl1*) and*RNA binding Fox-1 homolog 1* (*Rbfox1*) are differentially spliced but non-differentially expressed across spaceflight

AS is under the regulatory control of RNA binding proteins (RBPs) (Nilsen and Graveley, 2010). As such, it has long been hypothesized that DGE of these RBPs invokes downstream changes in AS. However, despite identifying numerous spaceflight-induced AS events, there was no evidence of spaceflight-induced DGE of RBPs (Supplemental Table 4). Surprisingly, instead, we found evidence of functionally significant AS of RBPs themselves.

Mbnl1 is an RBP with two tandem RNA binding domains (ZF1-2 and ZF 3-4) that acts as a splicing regulator of various musculoskeletal genes. *Mbnl1* undergoes AS of exon 2 that results in the generation of two isoforms, each with a different translational start codon (Figure 7A). In the exon 2-included isoform, the start codon resides in exon 2 and the final protein isoform contains both tandem binding domains. However, in the exon 2-excluded isoform, the start codon resides in exon 3 and the final protein isoform contains only the ZF 3-4 tandem RNA binding domain (Figure 7B). Importantly, these tandem RNA binding domains are not functionally equivalent; the ZF1-2 domain exhibits more motif specificity and five times more activity than the ZF3-4 domain (Lee, 2013). Therefore, it is interesting to note that the exon 2-included isoform, which contains both the ZF1-2 and ZF 3-4 domains, is 13% less abundant in the quadriceps of adult mice exposed to microgravity. In the gastrocnemius of mice exposed to spaceflight, the exon 2 included isoform was 3% less abundant (Figure 7C), a trend that did not reach statistical significance.

**Figure 7.**
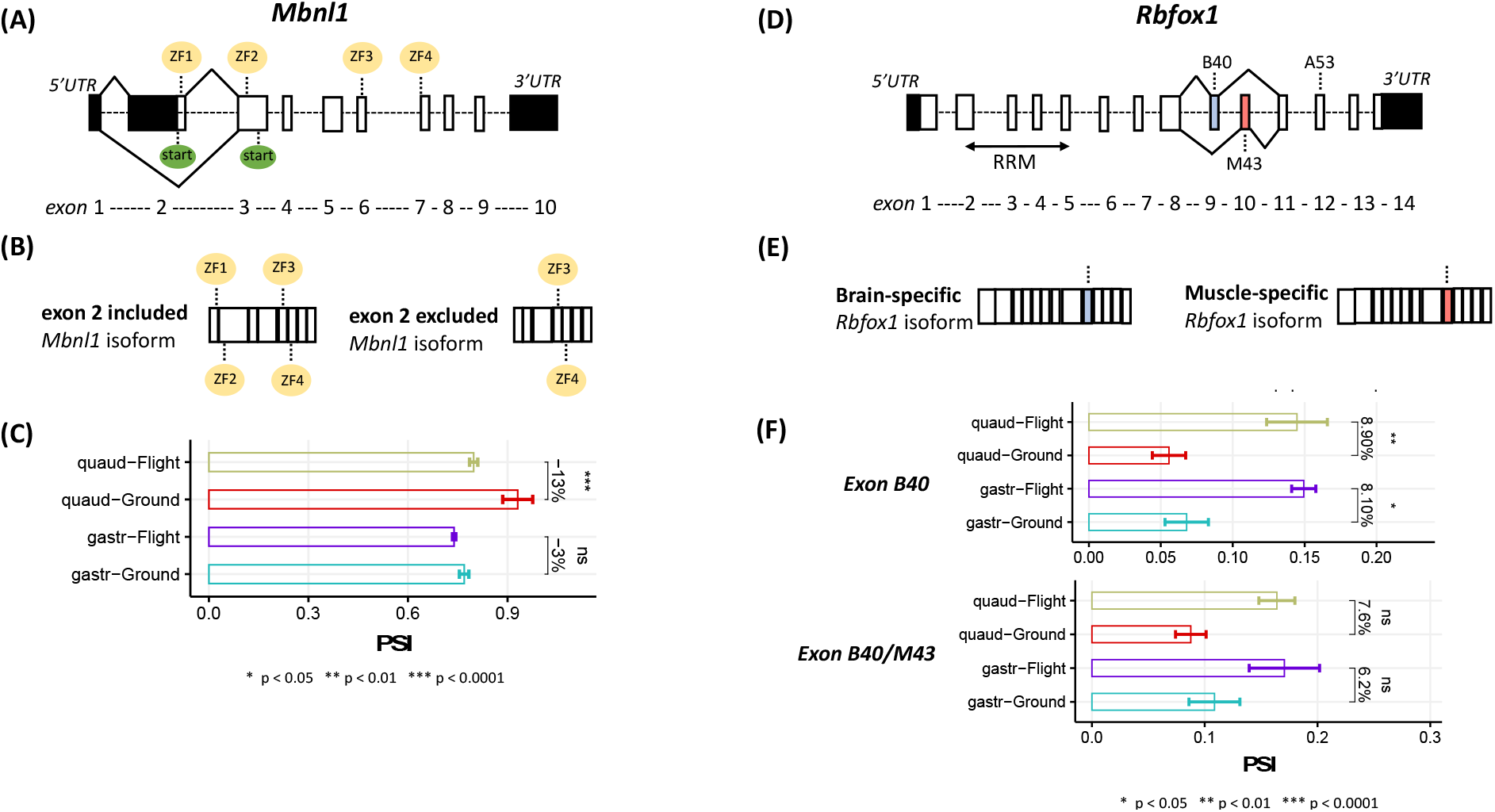
**(A)** Gene structure representation of the *Mbnl1* gene. Bars and dashed lines represent exons and introns, respectively, with exon numbering below. Untranslated regions (UTR) are denoted in black. Solid lines connect exon 2 to nearby splice junctions. Two start codons residing in exons 2 and 3 are depicted in green. Four zinc-finger domains (ZF1-4) residing in exons 2, 3, 6, and 7 are depicted in yellow. **(B)** Protein structure representation of the two *Mbnl1* isoforms. The exon 2 included *Mbnl1* isoform has exon 2 included and all four zinc finger domains (ZF1-4) are translated. The exon 2 excluded *Mbnl1*isoform has exon 2 excluded and only two zinc finger domains (ZF3-4) are translated. **(C)** Bar plot depicting the PSI (percent spliced in) value for exon 2 in ground and flight for quadriceps (top) and gastrocnemius (bottom). The delta PSI value indicates the difference in percent inclusion of exon 2 between flight and ground. Asterisks represent significance by two-tailed t-test **(D)** Gene structure representation of the *Rbfox1* gene. Bars and dashed lines represent exons and introns, respectively, with exon numbering below. Untranslated regions (UTR) are denoted in black. Solid lines connecting exons B40 (depicted in light blue) and exon M43 (depicted in red) to nearby splice junctions represent a mutually exclusive splicing. The RRM protein domain extends from exon 2 to 5 as depicted by the double sided arrow labeled as such. **(E)** Protein structure representation of the two *Rbfox1* isoforms. The brain-specific *Rbfox1* isoform has exon B40 included and exon M43 excluded while the muscle-specific *Rbfox1* isoform has exon B40 excluded and exon M43 included. **(F)** Bar plot depicting the PSI (percent spliced in) value of the exon B40/M43 mutually exclusive exon (MXE) event in ground and flight for quadriceps (top) and gastrocnemius (bottom). The delta PSI value indicates the difference in percent inclusion of exon B40/M43 between flight and ground. Asterisks represent significance by two-tailed t-test.

Rbfox1 is an RBP that acts as a splicing regulator in both muscle and brain tissue, with the tissue-specific function of this protein being regulated by a mutual exon exclusion (MXE) event involving B40 (brain-specific exon consisting of 40 base pairs) and M43 (muscle-specific exon consisting of 43 base pairs) (Figure 7D, 7E) (Conboy, 2017). While this has previously been annotated as a mutually exclusive event, we observed a significantly increased inclusion of B40 in both the in the quadriceps (8.9%) and gastrocnemius (8.1%) without a significant, reciprocal decrease in the inclusion of M43 in either muscle (Figure 7F).

## Discussion

While we set out first and foremost to characterize the role of AS in the physiological adaptation of the mouse hind limb postural and phasic muscles to microgravity, the work presented here has also provided novel insights more broadly into the reciprocal, tissue-specific relationship between DGE and AS. Co-transcriptional splicing was first documented as long as 30 years ago (Beyer and Osheim, 1988), but the molecular mechanisms underlying the coupling of DGE and AS have only recently been elucidated, including regulation of splicing by transcriptional elongation rate (Kadener, 2001) and modulation of splicing factor recruitment by nucleosome positioning and histone modifications (Spies, et al., 2009; Schwartz et al., 2009). In addition, more recent research has characterized a direct physical connection between RNA polymerase II and the spliceosome at the point of emergence of the pre-mRNA from the transcriptional machinery (Zhang et al., 2021). Together, this evidence suggests that these distinct mechanisms of transcriptome regulation (transcription and splicing) are intricately related. While it is believed that this intricate relationship is regulated by tissue-specific regulatory factors, DGE and AS have either been investigated together in a single biological system (cell type, tissue type, disease state, etc.) (Ullah et al., 2020; Brinegar et al., 2017) or separately across multiple biological systems (Shen-Orr et al., 2010; Yeo et al., 2004), without any comprehensive investigations of DGE and AS together across multiple biological systems. As such, the experiments here, documenting an equal but opposite propensity for the quadriceps and gastrocnemius to adapt their transcriptomes to microgravity via DGE or AS, respectively (see Figure 2A, 2B), provides the first evidence of a possible reciprocal, muscle-type-specific relationship between DGE and AS, such that each muscle preferentially employs one mechanism of transcriptome regulation at the other mechanism’s expense.

We hypothesize that the muscle-type-specific regulatory biases we identified are indicative of differences in energy availability across each muscle. While it has long been proposed that energy availability influences patterns of transcriptome regulation, only recently has it been shown that patterns of both DGE and AS vary across high and low ATP environments (Guantes et al., 2015) as a result of diminished supply of mitochondrial adenine nucleotides (AdNs: ATP, ADP, AMP) in atrophic skeletal muscle (Ji et al., 2019; Miller et al., 2020). In accordance with previous research (Roy et al., 1996; Recktenwald et al., 1999), at the level of muscle fiber immunostaining we observed more significant microgravity-induced atrophy in the quadriceps compared to the gastrocnemius (see Figure 1C), with the resistance to microgravity-induced atrophy in the gastrocnemius attributable to its more prominent slow-to-fast fiber type transition (see Figure 1A, 1B and 1D) (Fitts et al., 2001). Diminished adenine nucleotide supply in the more atrophied quadriceps may direct the use of AS, a less energy-dependent mode of gene expression, over DGE, a more-energy dependent mode of gene expression (Lynch and Marinov, 2015; Lane and Martin, 2010). In contrast, adequate supply of adenine nucleotides in the less atrophied gastrocnemius favors DGE over AS. While our results are restricted to a perturbation in gravity, a similar transcriptomic bias towards DGE over AS during development has been observed in the mouse gastrocnemius in utero (Brinegar et al., 2017), suggesting that our findings in the context of microgravity may reflect those of a “floating” amniotic environment.

Regardless of the mechanism of transcriptome regulation was preferentially employed in each muscle, we found a more significant role for AS than DGE in supporting microgravity-induced fiber type transitions. For example, in the absence of considerable DGE of genes encoding structural proteins in the muscle sarcomere, the increased reliance on fast twitch fiber function (whether in the 100% fast twitch quadriceps or the markedly slow-to-fast twitch transitioned gastrocnemius) in a microgravity environment was principally supported by extensive changes in AS of *Troponin T* (*Tnnt*). Our results provide evidence of three concomitant microgravity-induced isoform transitions, i) a move from *Tnnt3* pre-mRNA exon 17-to-16 inclusion (see Figure 3), ii) a *Tnnt3* acidic-to-basic residue change (see Figure 4), and iii) a *Tnnt1* lower-to-higher molecular weight change (see Figure 4). Taken in the context of previous research (Wang and Jin, 1997; Biesiadecki et al., 2007; Wei and Jin, 2011; Briggs et al., 1996; Ogut et al., 1999), when translated such isoforms would possesses N- and C-terminal alterations that, together, contribute in the three dimensional protein structures that strongly favor fast-twitch troponin I and tropomyosin binding as well as possess increased calcium sensitivity to contraction required by those fast-twitch fibers; that being said, it is unlikely that the aberrant splicing of *Troponin T* alone significantly enhanced the fast twitch function of either the quadriceps or gastrocnemius. In addition to the annotated events in *Tnnt3* and *Tnnt1*, there is evidence to suggest that AS of *Neb* and *Ryr1* may contribute to the function of fast twitch muscle. Specifically, the human homologs of *Neb* exon 127 (*NEB* exon 143) and *Ryr1* exon 70 (*RYR1* exon 70) are known to be preferentially included in the mRNA of fast twitch muscles and reciprocally excluded in slow twitch muscles (Tang et al., 2015; Lam et al., 2018). While the mechanism that contributes to increased use of these isoforms in fast-twitch muscle remains unknown, the microgravity-induced inclusion of *Neb* exon 127 and *Ryr1* exon 70 can be taken as further evidence of splicing-supported fiber type transitions in response to extended spaceflight.

Similar to microgravity-induced fiber type transitions, we found a more significant role for AS than DGE in supporting microgravity-induced muscle atrophy. Unlike the role of AS in the slow-to-fast fiber transition, the precipitation of muscle atrophy via AS is not an entirely novel concept; most support for this phenomenon comes in the context of atrophic musculoskeletal diseases. For example, the aberrant splicing of *NEB* exon 143 and *RYR1* exon 70, have been shown to be required for the development of atrophic musculoskeletal diseases in humans (Garg et al., 2017; Kimura et al., 2005). While we observed aberrant splicing of both *Neb* exon 127 and *Rry1* exon 70 in the muscle transcriptome of our mice, those changes opposed the AS trends that would generate atrophy. As such, we found no evidence to support a “canonical” role of AS in precipitating muscle atrophy. Despite this, we were successful in identifying a relatively novel mechanism of AS-induced atrophy: the regulation of large musculoskeletal gene transcript length. The unique composition of musculoskeletal tissue in sarcomere units necessitates the employment of some of the largest proteins in nature. This includes titin and nebulin (see Figure 6). Owing to their size (350 exons and 166 exons, respectively) and complexity, the pre-mRNAs of these genes lend themselves to extensive regulation via AS. In fact, AS is well-characterized in regulating developmental changes in the size of these proteins; for example, on average, the size of titin decreases by approximately 7.0% during musculoskeletal development as a result of extensive AS within the PEVK domain (Ottenheijm er al., 2009). In contrast to the typical developmental shortening of the *Ttn* transcript, we characterized AS-directed extension of the *Ttn* transcript as a result of extended exposure to microgravity, which, together with the AS-directed shortening of the *Neb* transcript, serves to diminish the overall contractile strength of the muscle sarcomere. In addition, differences in these AS events across the quadriceps and gastrocnemius may help to further explain the resistance of postural muscles to microgravity-induced atrophy, as the gastrocnemius displayed AS-directed shortening of the *Neb* transcript but no AS-directed extension of the *Ttn* transcript.

In the absence of significant differential expression of RBPs in either the of the postural or phasic limb muscles studied here, we discovered functionally significant spaceflight-induced AS of RBPs themselves, a likely upstream provocation that can account for changes in AS of downstream gene targets. For example, *Mbnl1* was found to be significantly alternatively spliced in the quadriceps in microgravity and *Rbfox1* significantly alternatively spliced in both the quadriceps and gastrocnemius in microgravity (see Figure 7). Mbnl1 regulates the splicing patterns of various musculoskeletal genes, including *Tnnt1*, *Tnnt3*, *Ttn*, and *Ryr1*. Specifically, Mbnl1 has been motif-mapped to the 3’ variable region of *Tnnt3*, exon 5 of *Tnnt1*, the PEVK domain of *Ttn*, and exon 70 of *Ryr1* (Lee, 2013), all events that were found in the work presented here to be significantly alternatively spliced as a result of exposure to microgravity (see Figure 3, 4, 5, 6). Therefore, AS of *Tnnt1*, *Tnnt3*, *Ttn*, and *Ryr1* may be the result of spaceflight-induced exclusion of the functionally superior ZF1-2 domain of *Mbnl1*. Similar to Mbnl1, Rbfox1 regulates the splicing patterns of various musculoskeletal genes, including *Ryr1* and *Mybpc1*. Rbfox1 has been motif-mapped to exon 70 of *Ryr1* and exon 3 of *Mybpc1* (Pedrott et al., 2015), both events that were found in the work presented here to be significantly alternatively spliced as a result of exposure to microgravity (see Figure 5, Supplemental Tables 5 and 6). Therefore, AS of *Ryr1* and *Mybpc1* may be the result of spaceflight-induced inclusion of the aberrant brain-specific exon in place of the muscle-specific exon of *Rbfox1* mRNA. Taken together, these data suggest that pre-mRNA targets for differential AS are subject to control by AS of the RBP splicing factors itself that ultimately directs AS of that target pre-mRNA.

While our experiments employed adult mice (32 weeks), it is unlikely that age alone can account for our findings. Previous characterizations of the musculoskeletal adaptation to spaceflight have occurred in the context of mice ranging from 8-20 weeks at the time of spaceflight (Harrison et al., 2003; Gambara et al., 2017; Camerino et al., 2013; Chakraborty et al., 2020), requiring inquiry as to the role age may have played in generating the phenotypes we observed and the transcriptomic alterations we characterized.

First, there is evidence in humans to suggest that a fast-to-slow fiber type transition accompanies aging (Ohlendiek, 2011). Therefore, the presence of a microgravity-induced slow-to-fast fiber transition suggests that, despite the opposing force of aging on fiber type composition, the slow-to-fast fiber plasticity of skeletal muscle fibers is maintained even at an increased age. In contrast to fiber transitions, aging mirrors the effect of microgravity on muscle atrophy (Ohlendiek, 2011); it is possible that the advanced age of our mice compared to early age mice in other studies (Harrison et al., 2003; Gambara et al., 2017; Camerino et al., 2013; Chakraborty et al., 2020) amplified the microgravity-induced atrophy that was observed here. Most important to our work, however, is the relationship between modes of transcriptomic regulation and age. While relative employment of DGE and AS fluctuates with age and organismal development, most of this fluctuation occurs immediately preceding, at, and immediately following birth (until approximately post-natal day 28; Brinegar et al., 2017). In addition, there is evidence in both mice and humans to suggest relative stability of the striated muscle transcriptome across adult years and even into later life (Balliu et al., 2019). Therefore, it is unlikely that the transcriptome alterations we characterized are simply a product of the age of the mice employed. In fact, investigations of microgravity-induced differential gene expression in younger mice (Gambara et al., 2017; Chakraborty et al., 2020) showed downregulation of metabolic and mitochondrial pathways similar to what we identified in our older mice. Therefore, while AS has not been adequately investigated in the context of spaceflight, it can be speculated that extensive microgravity-induced AS would be identified regardless of age.

In summary, we have shown that 1) AS and DGE vary in a reciprocal manner, possibly in response to tissue-specific energy availability; 2) AS plays a more functionally significant role than DGE in supporting microgravity-induced phenotypes; 3) AS augments fast twitch fiber function via aberrant splicing of *Tnnt3*, *Tnnt1*, *Neb*, and *Ryr1*; 4) AS contributes to muscle atrophy via splicing-directed shortening of *Neb* and extension of the *Ttn*; and 5) RBPs, transregulators of AS, are themselves differentially spliced while being non-differentially expressed. Together, the results of our work emphasize the importance of AS in determining the plasticity and functional status of the skeletal muscle transcriptome. Considering the current development of novel treatment methods for atrophic musculoskeletal diseases using small-molecule splicing modulators (Jarecki et al., 2005; Cheung et al., 2018; Ando et al., 2020), it is possible that further characterization of microgravity-induced AS targets will allow for the development of small-molecule splicing modulator-based therapies to prevent microgravity-induced muscle atrophy and other detrimental physiologic adaptations to spaceflight.

## Materials and Methods

### Animals

All the animal procedures were performed according to the guidelines of the Chancellor’s Animal Research Committee at UCLA (Protocol # 2009-127) and NASA Institutional Animal Care and Use Committees. Twenty 32-week-old female BALB/c mice (Taconic Biosciences, NY) were used in this study. All mice were housed at Kennedy Space Center (KSC) for 4 weeks before rocket launch and were randomly divided into ground control group and flight groups (n=10 per group). On June 3^rd^, 2017, mice of in the flight group were transported to the International Space Station (ISS) as part of SpaceX CRS-11 and kept at ISS for the full 9 weeks of experimentation, while mice of ground control group were kept at KSC for the same duration. The mice housed at KSC were in the NASA’s Rodent Research Hardware System and were fed food bar diet ad libitum. The food bar essentially has added moisture in the standard rodent chow to form a bar form and prevent it from floating in microgravity.

### Sample preparation

At the end of the study, all mice were euthanized at ISS or KSC by our NASA partners on site. Mice were then were deeply frozen in situ under −80c or colder until later transportation to UCLA on dry ice. Frozen carcasses were thawed on wet ice before tissue dissection. Mice skeletal muscles were dissected separately and either fixed in 10% neutral buffered formalin or preserved in *RNAlater* solution.

### Immunohistochemistry and morphological analysis

Formalin-fixed paraffin-embedded sections of mouse muscle (5um thickness) from gastrocnemius and quadriceps were cut with a microtome and mounted on charged slides. Sections were either subjected to standard hematoxylin-eosin (H&E) staining for overview, or immunolabeled with the following anti-myosin heavy chain (MyHC) isoform antibodies: type I MyHC isoform (Abcam, Cat# ab11083); type II MyHC isoform (Abcam, Cat# ab51263). The sections were co-stained with an anti-laminin antibody (Abcam, Cat# ab11575) to allow measurement of fiber size. In all protocols, donkey anti rabbit Alexa-488 conjugated secondary antibody was used for laminin staining, donkey anti-mouse Alexa-594 conjugated secondary antibody for type I MyHC antigen staining, and anti-mouse Alexa-488 conjugated secondary antibody for type II MyHC antigen staining. Photomicrographs were acquired using Olympus BX 51 and IX 71 microscopes equipped with Cell Sense digital imaging system (Olympus, Japan). To assess the cross-sectional area (CSA) of the different myofiber types, digitized photographs were acquired and the myofiber CSA was automatically measured by means of ImageJ 1.45 g (NIH, freeware imaging software).

### RNA extraction and purification

Total RNA was isolated from mouse gastrocnemius (n=3) and quadriceps (n=3) of each experimental group (flight and ground control) using the acid guanidinium thiocyanate-phenol-chloroform extraction followed by silica-membrane purification. Briefly, frozen tissue samples were minced into small pieces (1mm x 1mm x 1mm). A homogeneous lysate was achieved by adding lysing buffer and gentle ultrasound vibration on ice. Tissue lysate was centrifuged and the supernatant was used for RNA phenol/chloroform extraction. After phase separation, the aqueous layer was transferred and mixed with an equal volume of 70% ethanol. Total RNA was then extracted using Rneasy spin columns from the Rneasy micro Kit (Qiagen, Hilden, Germany) according to the manufacturer’s protocol.

### RNA seq analysis

For each replicate RNA sample, a cDNA library was prepared and sequenced by the UCLA Technology Center for Genomics and Bioinformatics using an Illumina sequencer, generating an average of 41.5 million single-end 50bp reads. The resulting RNASeq reads were aligned to the mouse genome (mm10) reference using STAR software (Dobin et al., 2013). Quality of the RNA-seq dataset was confirmed by read depth and mapping statistics (Supplemental Figure 1A). Read depth for all samples was at or above 30 million uniquely mapped reads (the desired read depth for differential splicing analyses), with the exception of one quadricep ground sample with 27.4 million uniquely mapped reads. The uniquely mapped read percentage for all samples was above 80%, the desired mapping percentage for differential splicing analyses. Transcript abundance was measured directly from FASTQ files as TPM (Transcripts Per Million) using kallisto (Bray et al., 2016) and summarized into gene expression matrix by R package ‘tximport’ (Soneson et al., 2015). Lowly expressed genes (TPM <= 5 in all samples) were filtered out before conducting differential analysis with DeSeq2 (Love et al., 2014). For each comparison, genes with an absolute log2 fold change > log2(1.5) and an FDR (false discovery rate)-adjusted p-value < 0.01 were assumed to be differentially expressed genes.

Alternative splicing events were detected and quantified as Percent Spliced In (PSI) values by rMATS-turbo (Shen et al., 2014) using junction reads (reads spanning the splicing junctions). AS events with low junction read support (≤ 10 average junction reads, ≤ 10 total inclusion junction reads or ≤10 total skipping junction reads over all 12 samples), or with extreme PSI value ranges (PSI ≤ 0.05 or ≥ 0.95 in all 12 samples) were excluded from downstream analysis. Differential splicing analysis was then performed using rMATS-turbo by comparing the replicate ground samples and flight samples in each tissue. Differential splicing events were identified by the following criteria: 1) >10 average junction reads (inclusion and skipping junction reads) in both groups; 2) no extreme PSI values (PSI ≤ 0.05 or PSI ≥ 0.95 for all 6 samples in the comparison); 3) FDR < 0.01; and 4) absolute change in PSI (|ΔPSI|) > 0.05. Five AS event types were annotated (Supplemental Figure 1B), including skipped exon (SE), alternative 5’ splice site (A5SS), alternative 3’ splice site (A3SS), mutually exclusive exon (MXE), and retained intron (RI). SE and MXE, AS event types focused on in this work, composed approximately 50% and 20% of all AS events, respectively. Gene ontology (GO) analysis was performed to reveal enriched functional pathways affected by significant gene expression changes as well as alternative splicing changes using EnrichR (Chen et al., 2013; Kuleshov et al., 2016; Xie et al., 2021). The top enriched GO terms with smallest adjusted p values were visualized as bar plots.

---

**Supplemental Figure 1.**
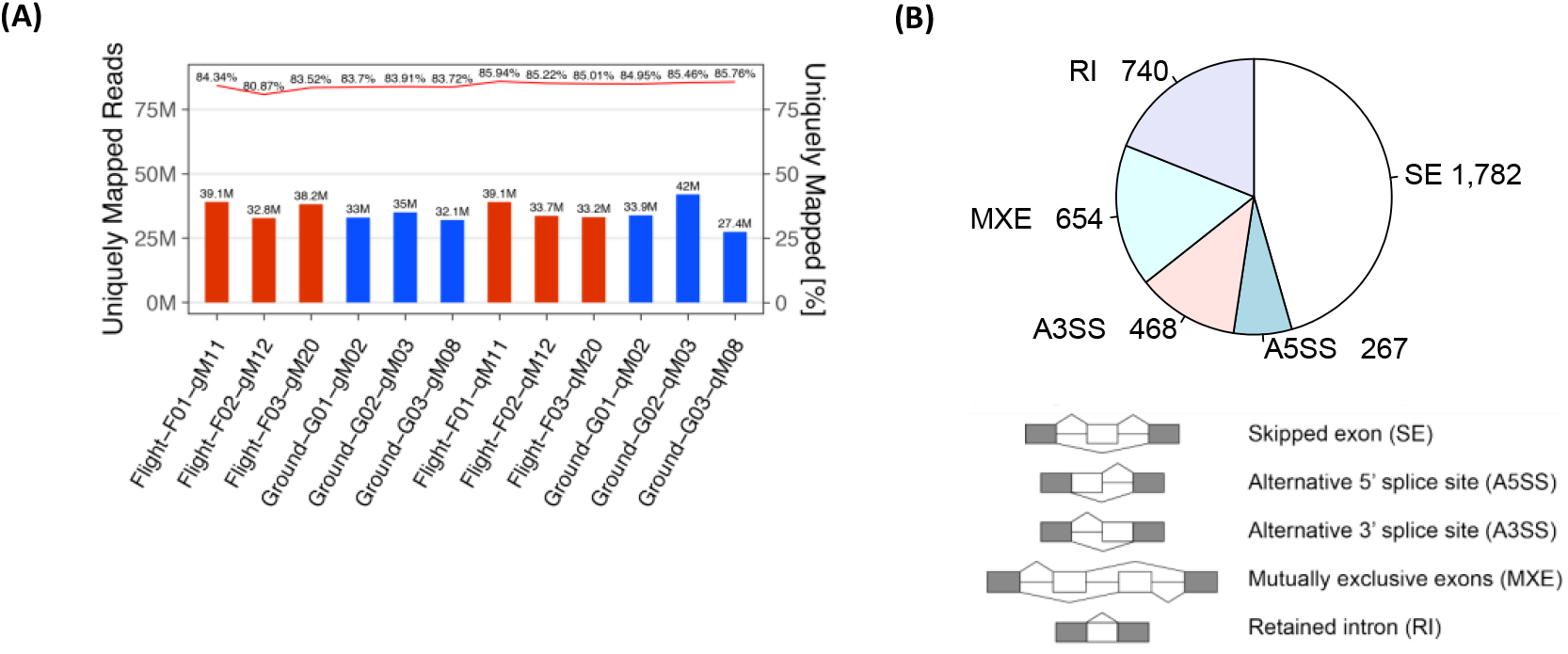
**(A)** Summary of read depth and mapping statistics for RNA-seq dataset. **(B)** Summary table of AS events detected by rMATS-turbo after filtering by read coverage and PSI value range. SE, skipped exon; A5SS, alternative 5’ splice site; A3SS, alternative 3’ splice site; MXE, mutually exclusive exon; RI, retained intron. Representative images below depict examples of the above listed alternative splicing events.

**Supplemental Table 1.** Enrichment by gene ontology of upregulated genes in gastrocnemius and quadriceps. Reported here are the GO term (top five for each category), GO category, and adjusted p-value.

|  | *Gastrocnemius* |  |  | *Quadriceps* |  |  |
| --- | --- | --- | --- | --- | --- | --- |
|  | GO Term | GO category | Adjusted P-value | GO Term | GO category | Adjusted P-value |
| Upregulated Genes | Spinal cord motor neuron differentiation | Biological Process | 0.0821 | Synaptic transmission, glycinergic | Biological Process | 0.03089 |
|  | Cardiac ventricle formation |  | 0.0821 | Inner ear receptor differentiation |  | 0.03089 |
|  | Lymphoid progenitor cell |  | 0.0821 | Response to catecholamine |  | 0.03089 |
|  | Dendrite extension |  | 0.0821 | Cellular response to monoamine stimulus |  | 0.03089 |
|  | Cell differentiation in spinal cord |  | 0.0821 | Positive regulation of myeloid leukocyte cytokine production involved in immune response |  | 0.03089 |
|  | Peroxisome proliferator activated receptor binding | Molecular Function | 0.08081 | Glucocorticoid receptor binding | Molecular Function | 0.08182 |
|  | AT DNA binding |  | 0.08081 | Protein phosphatase 1 binding |  | 0.08182 |
|  | Retinoid X receptor binding |  | 0.08081 | Glycine binding |  | 0.08182 |
|  | Filamin binding |  | 0.08081 | Inhibitory extracellular ligand-gated ion channel activity |  | 0.08182 |
|  | Sequence-specific double-stranded DNA binding |  | 0.08081 | Ligand-gated anion channel activity |  | 0.08182 |
|  | Synaptonemal structure | Cellular Component | 0.175 | Cytoplasmic microtubule | Cellular Component | 0.02628 |
|  | Synaptonemal complex |  | 0.175 | Microtubule |  | 0.1374 |
|  | Cytoplasmic microtubule |  | 0.175 | Polymeric cytoskeletal fiber |  | 0.1441 |
|  | Contractile actin filament binding |  | 0.175 | Microtubule cytoskeleton |  | 0.2399 |
|  | Stress fiber |  | 0.175 | Integral component of plasma membrane |  | 0.2733 |

**Supplemental Table 2.**
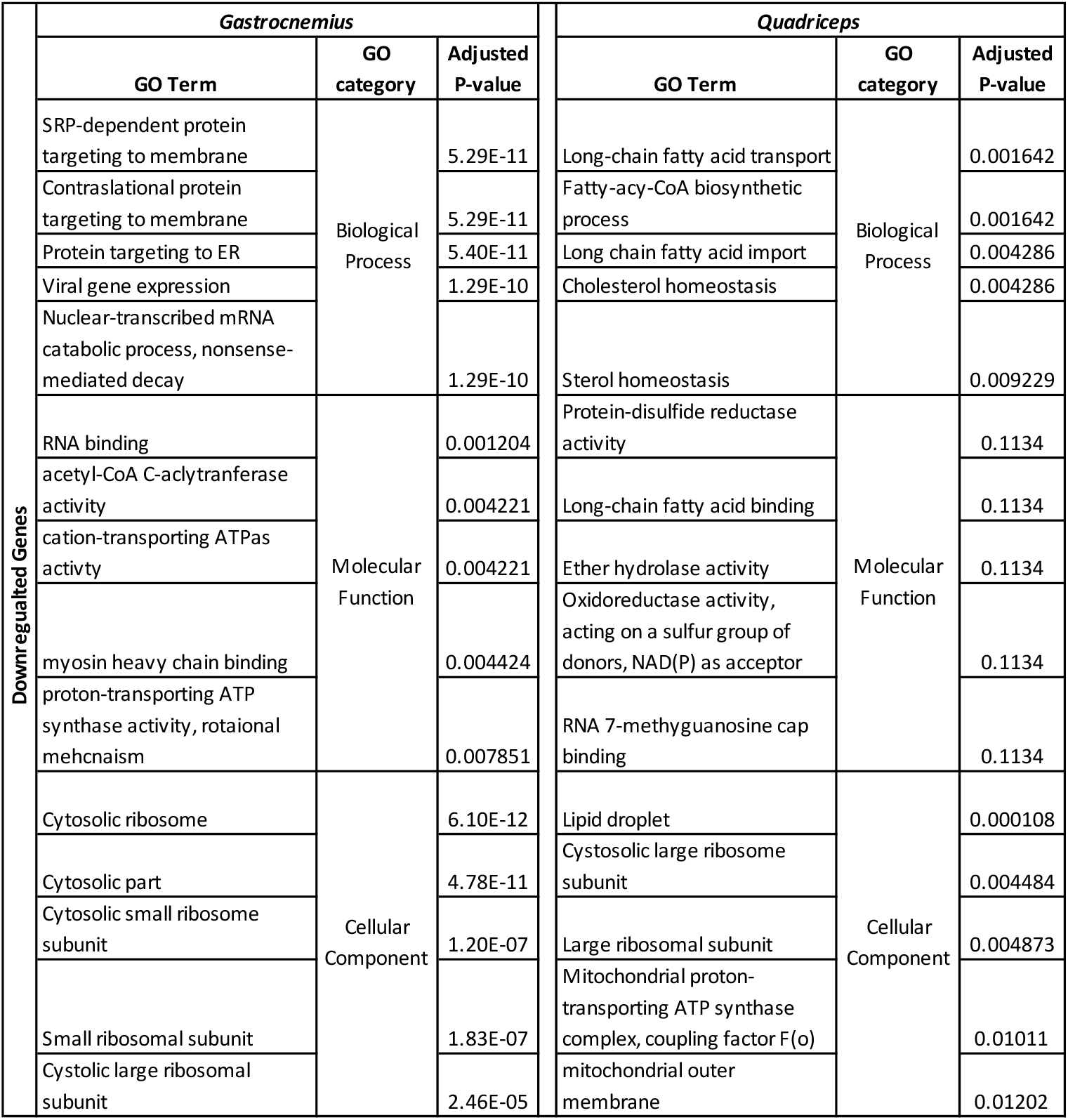
Enrichment by gene ontology of downregulated genes in gastrocnemius and quadriceps. Reported here are the GO term (top five for each category), GO category, and adjusted p-value.

**Supplemental Table 3.**
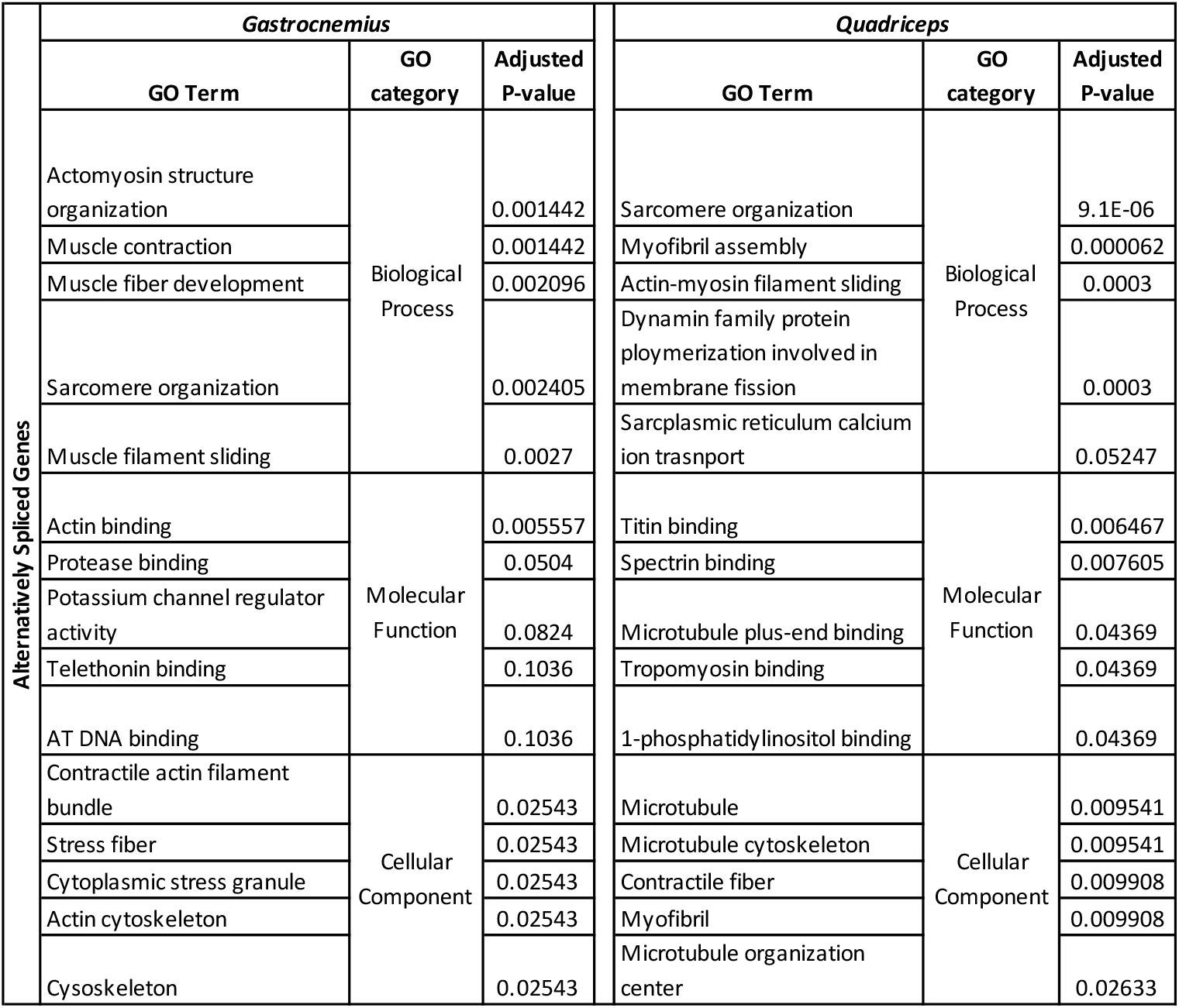
Enrichment by gene ontology of alternatively spliced genes in gastrocnemius and quadriceps. Reported here are the GO term (top five for each category), GO category, and adjusted p-value.

**Supplemental Table 4.**
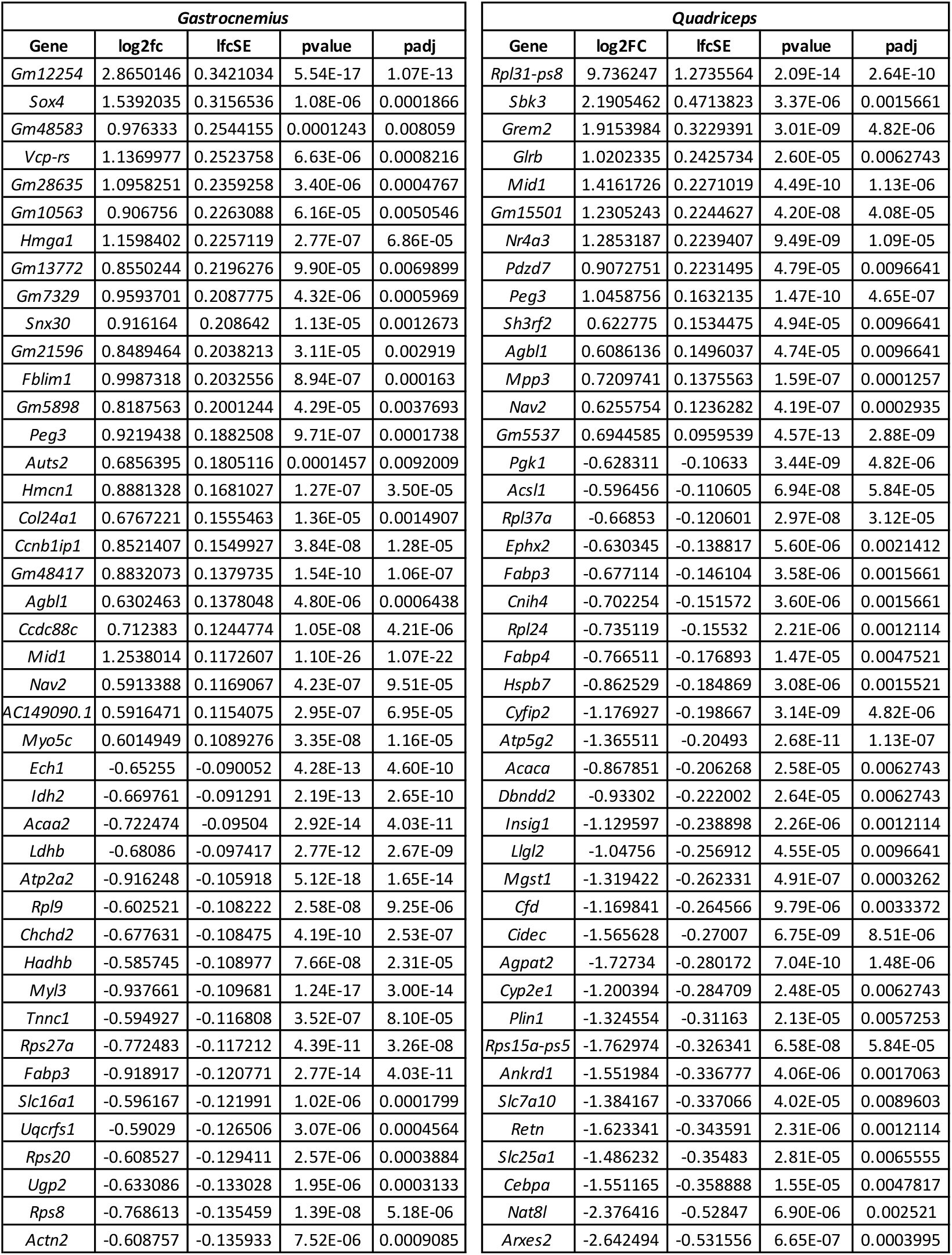
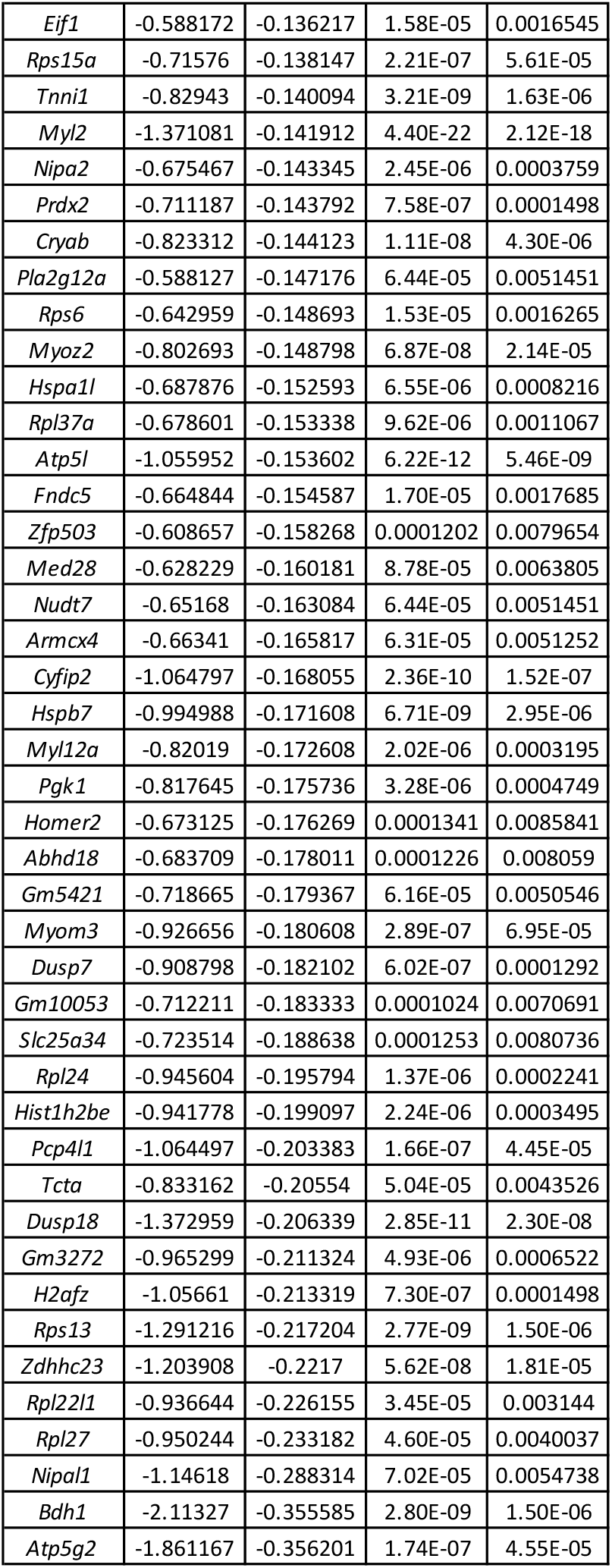
86 and 43 genes were significantly differentially expressed across spaceflight in the gastrocnemius and quadriceps, respectively. Reported here are the gene symbol, log2FC, p value, and FDR.

**Supplemental Table 5.**
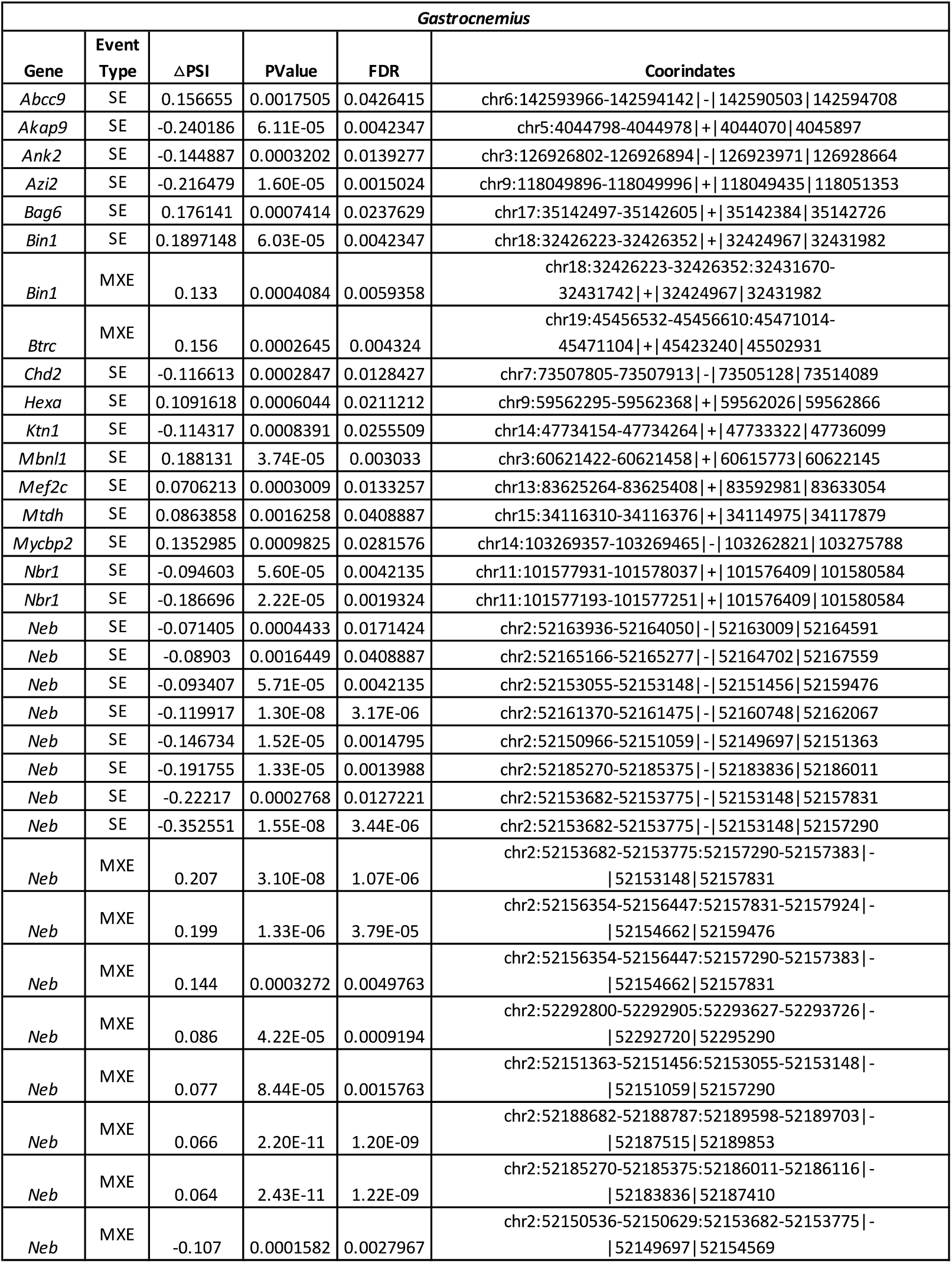
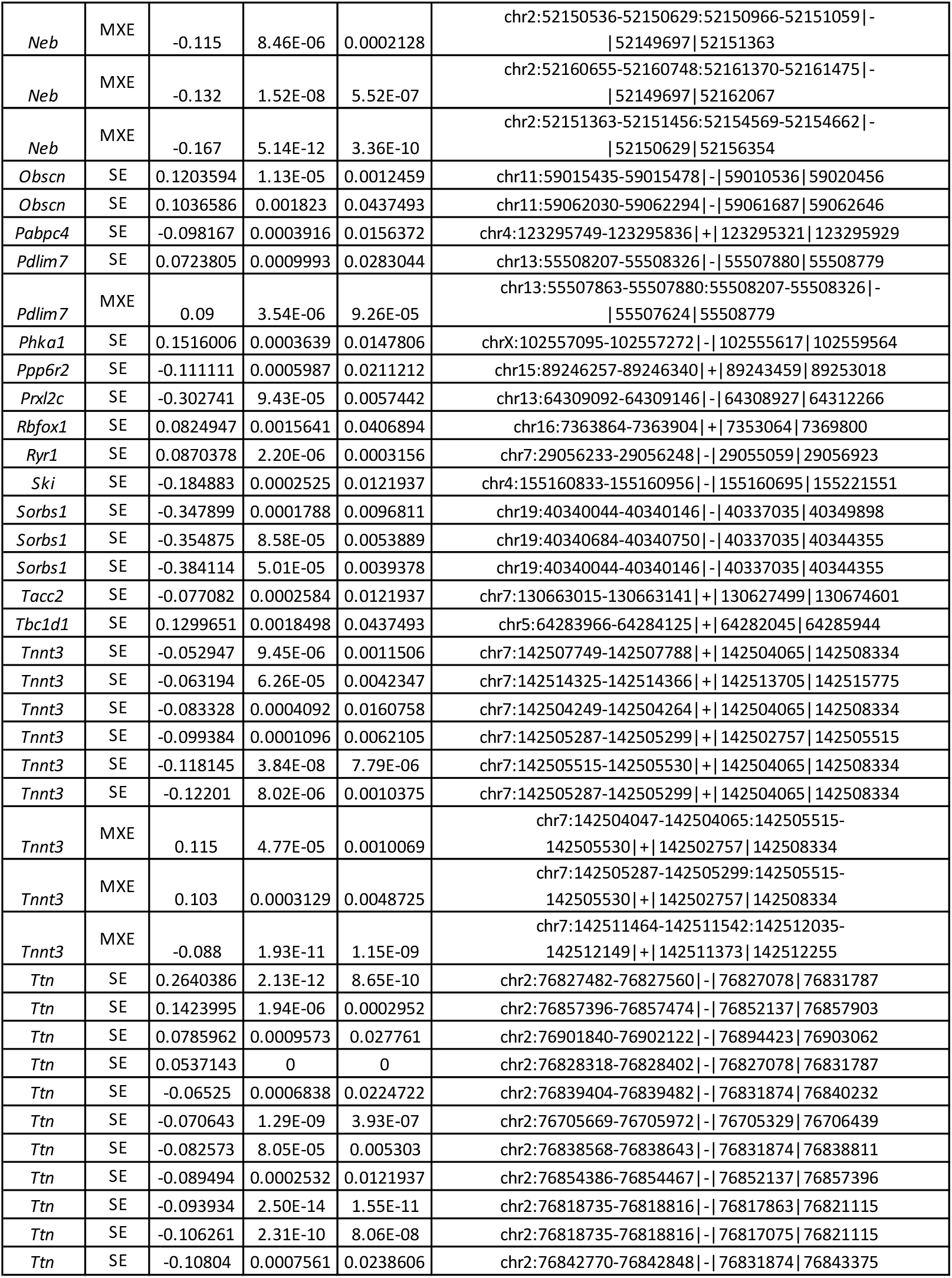
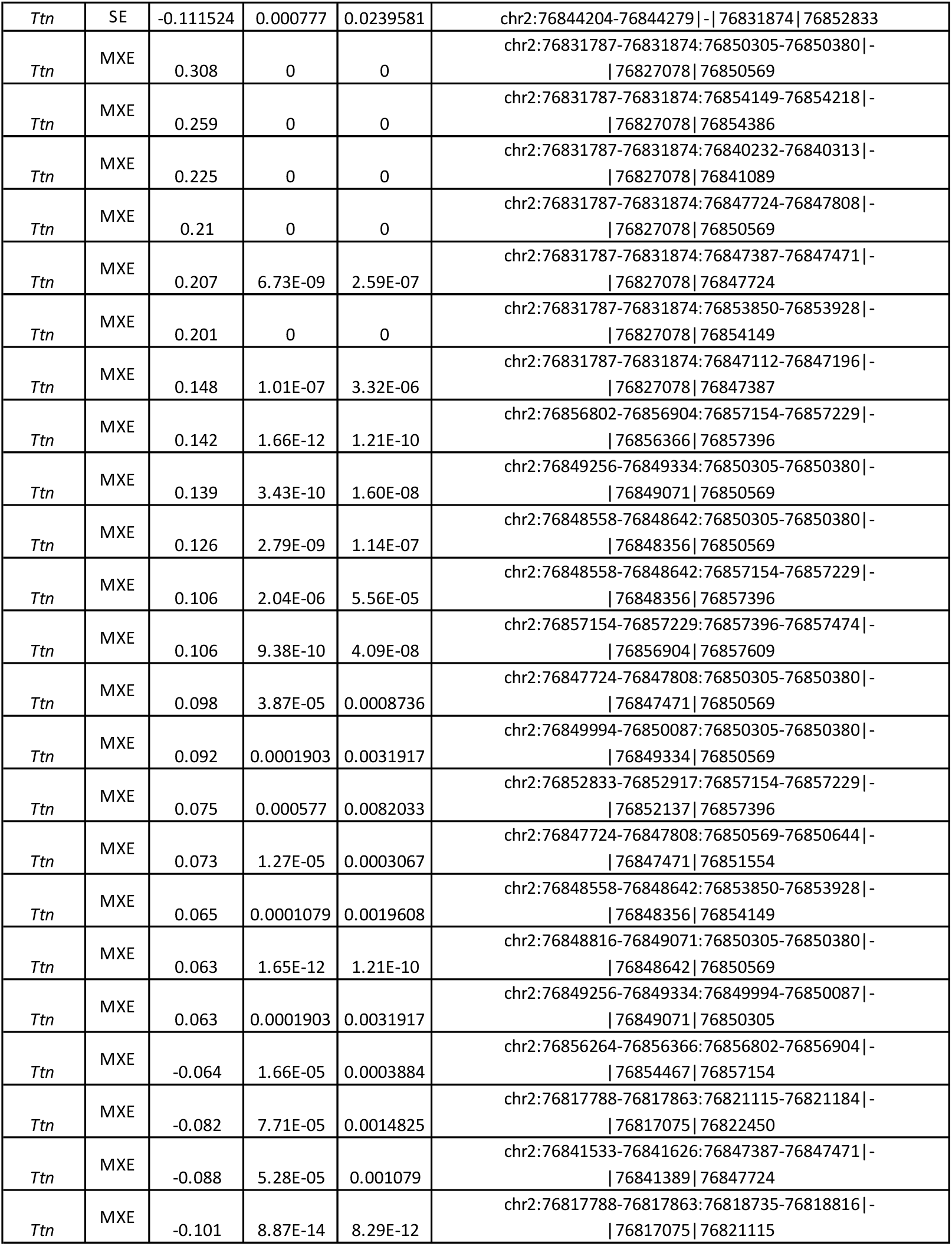
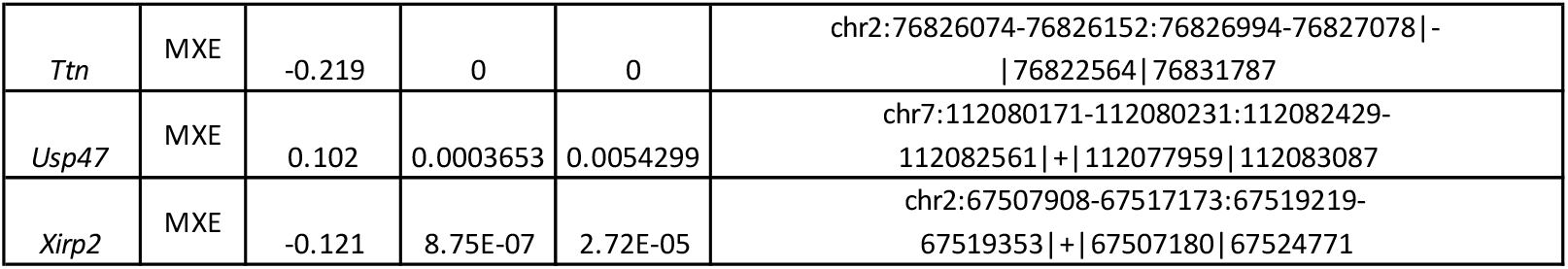
99 statistically significant alternative splicing events were identified across spaceflight in the gastrocnemius, representing 32 alternatively spliced genes, respectively. Reported here are the gene symbol, event type, delta PSI, p value, and FDR.

**Supplemental Table 6.**
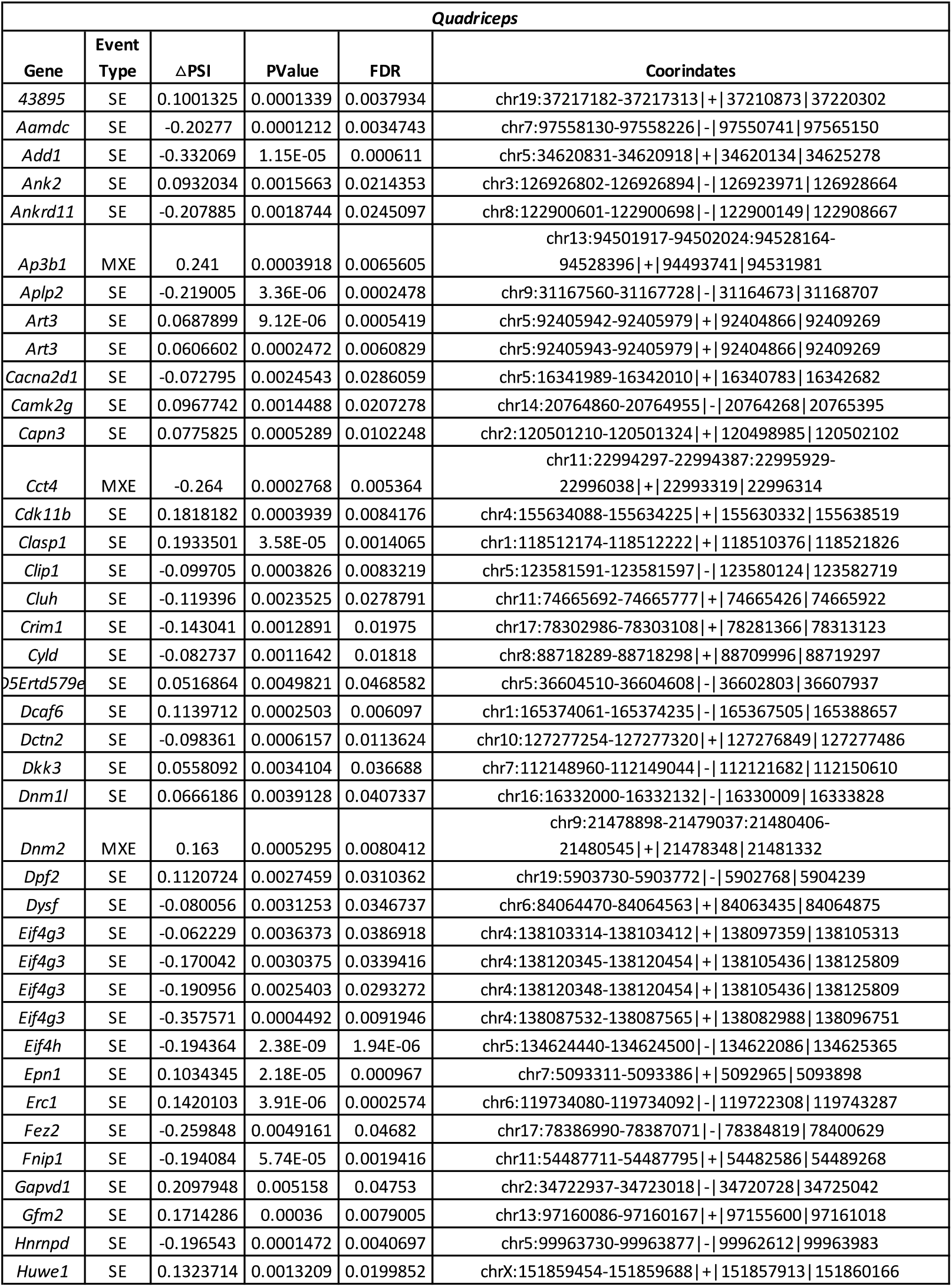
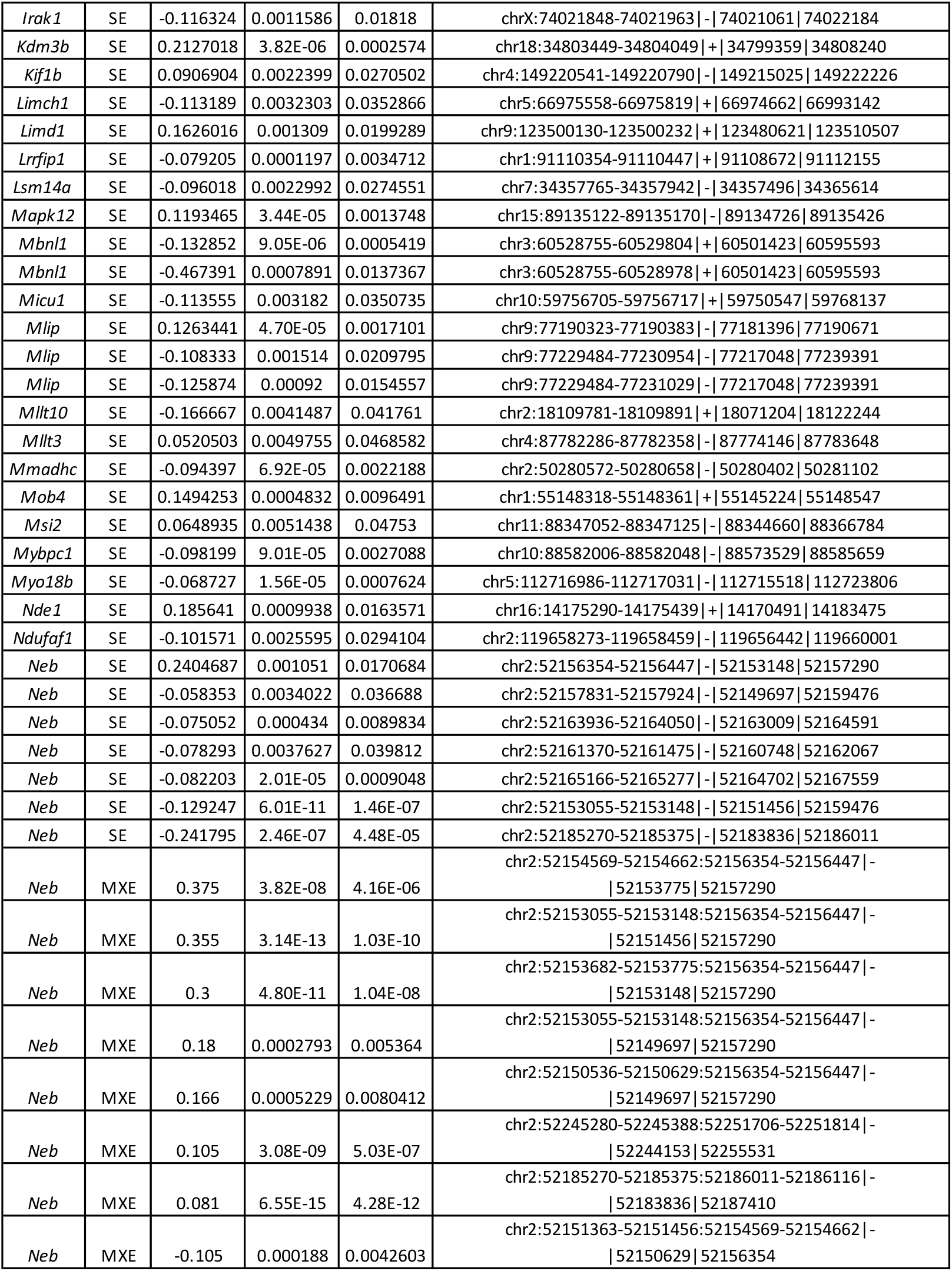
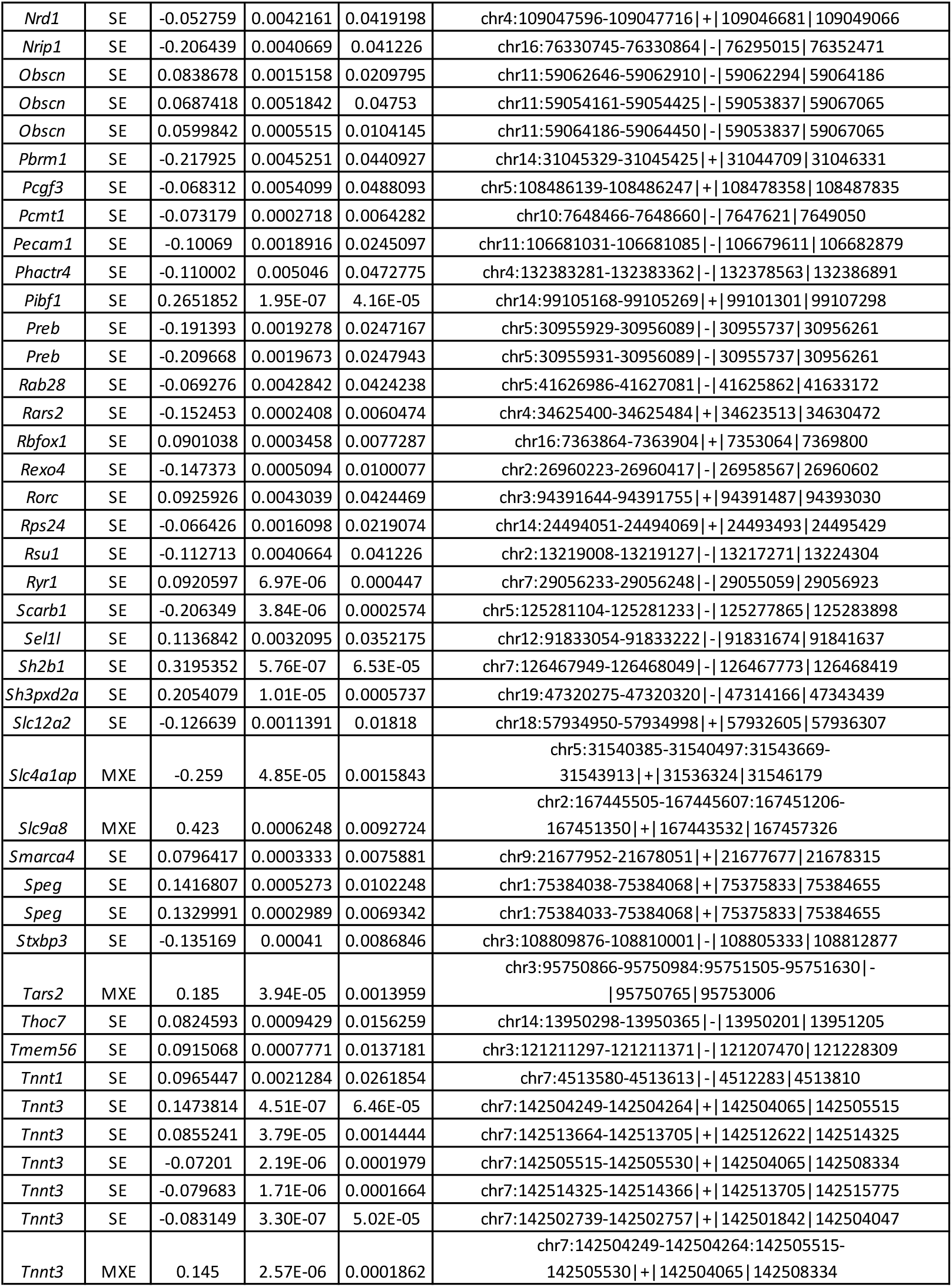
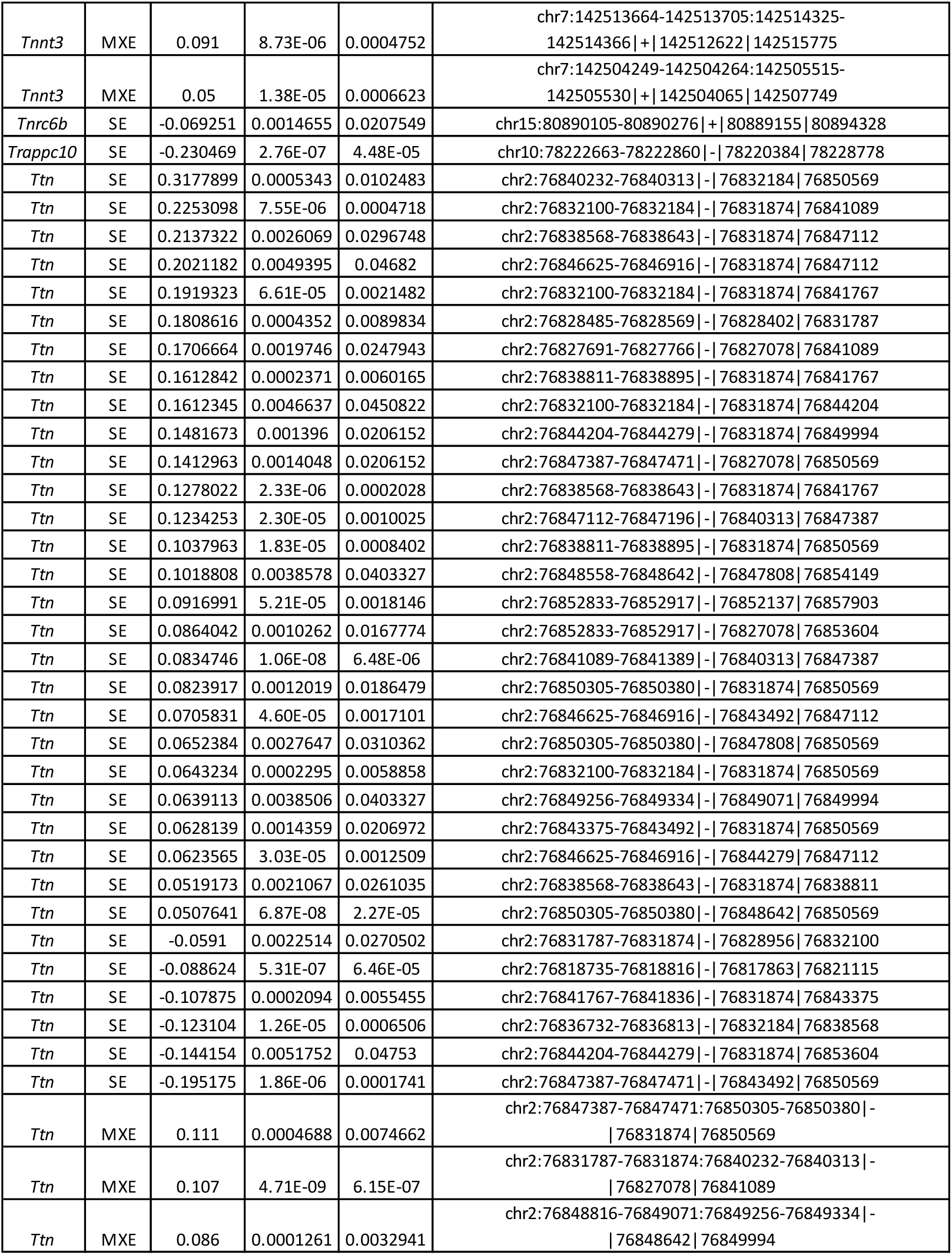
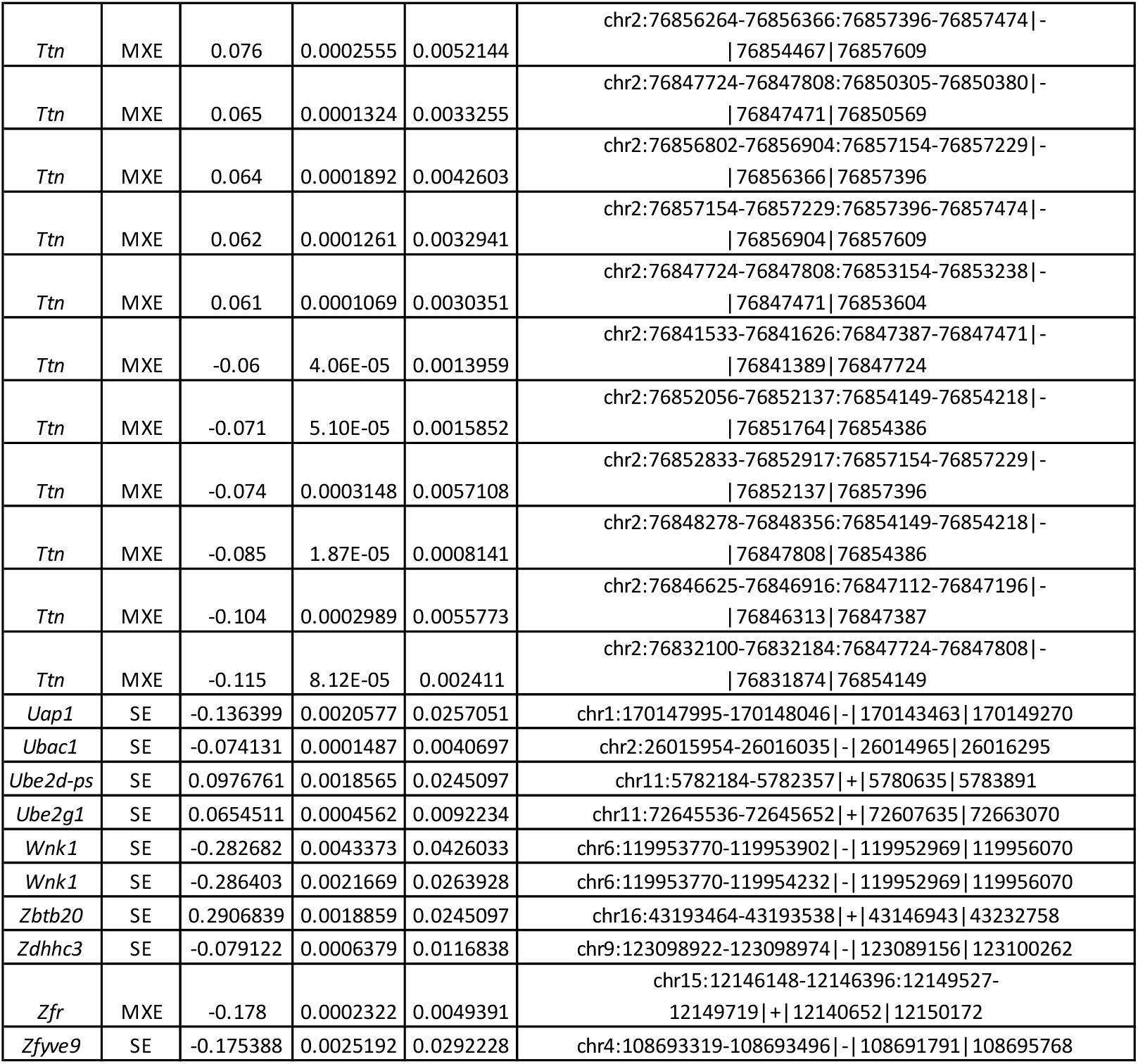
181 statistically significant alternative splicing events were identified across spaceflight in the quadriceps, representing 103 alternatively spliced genes, respectively. Reported here are the gene symbol, event type, delta PSI, p value, and FDR.

